# Identification and validation of liquid biopsy-based methylation biomarkers: a germ cell tumor subtype-specific study

**DOI:** 10.1101/2025.11.04.686501

**Authors:** Ferdinand W. Janssen, Ad J. M. Gillis, Lennart A. Kester, Hirokazu Takami, Koichi Ichimura, Thomas F. Eleveld, Leendert H.J. Looijenga

## Abstract

Human germ cell tumors (GCTs) occur in infants, children, and adults, and present as germinomatous and/or non-germinomatous (embryonal carcinoma, teratoma, yolk sac tumor (YST), and choriocarcinoma) histologies at gonadal or extragonadal locations. Accurate subtyping is crucial for prognosis and treatment, but current clinical biomarkers lack sensitivity and specificity (serum proteins) or require a tissue biopsy (immunohistochemistry). Hence, less-invasive and improved subtype-specific biomarkers have potential for clinical utility. We conducted a meta-analysis of DNA methylation data (450K/EPIC array) from 15 (three original and 12 published) datasets including 713 GCTs, 109 healthy testis, and 221 healthy peripheral blood samples, revealing that GCT histology is the primary driver of methylation profiles, regardless of tumor location and patient’s age or sex. Per subtype, we identified differentially methylated regions as potential biomarkers. As proof of concept, we identified and validated two YST-specific biomarkers, i.e., *APC* and *DPP7* promotor methylation, using methylation-sensitive restriction enzyme-based qPCR, of which *DPP7* was also detectable in GCT serum-derived cell-free DNA. In conclusion, we present a novel method for *in silico* identification and *in vitro* and *in vivo* validation of YST subtype-specific liquid biopsy-based biomarkers. Our bioinformatic pipeline is easily transferrable encouraging additional applications in pan(pediatric)-cancer studies beyond GCTs.

## Introduction

Human germ cell tumors (GCTs) are a heterogeneous group of neoplasms, derived from embryonic germ cells. They can be diagnosed in infants, children, and adults, and localize in the gonads (testis and ovary) and extragonadal sites, including the brain [1]. Although overall rare in the general population, testicular germ cell tumors (TGCTs) account for >90% of testicular tumors and are the most common cancer type in young men between 15 and 40 years of age [2, 3]. In contrast, 20-25% of ovarian tumors have a germ cell origin (70% in the first two decades of life), and only 2-3% of (pediatric) brain tumors are GCTs [4, 5]. Prognosis of GCT patients is generally good with a 5 year overall survival rate of 65-96% depending on defined risk groups [6-8]. Platinum-based chemotherapy resistance (5-10% of all patients) remains the most important predictor of poor outcome [9]. Overall, GCTs are classified into different types based on their origin, histology, (epi)genetic characteristics, and anatomical localization, of which type I and II are the most prevalent [1]. Type I GCTs originate from early embryonic germ cells and typically occur in neonates and children <6 years. They consist of benign teratoma (TE) or malignant yolk sac tumor (YST) (or a combination of both) and can be found along the midline of the body, from the sacrococcygeal region, the gonads (ovary and testis), and other extragonadal sites like the retroperitoneum, abdomen, mediastinum, and brain. In contrast, type II GCTs are malignant by definition and develop from more mature embryonic germ cells. They clinically manifest post-puberty, mostly in young adults, and typically localize in the testis. Type II GCTs are histologically and clinically grouped into germinomatous GCTs (gGCTs) and non-germinomatous GCTs (ngGCTs) [1]. gGCTs are called seminoma (SE) when occurring in the testis, dysgerminoma (DG) in the ovary, and germinoma (GER) in the brain. ngGCTs consist of embryonal carcinoma (EC), TE, YST, or choriocarcinoma (CHC), most commonly as mixed components, possibly combined with a gGCT component, while they can also (in rare cases) be histologically pure. Although type I and type II GCTs can share histology (TE and YST), their origin and development are distinct. Importantly, type II TGCTs always develop from a germ cell neoplasia *in situ* (GCNIS) precursor lesion and show a characteristic gain of the short arm of chromosome 12, while type I TGCTs typically do not, although additional copies of the 12p13 region can be present [1].

Correct identification of the GCT histological subtypes is clinically relevant as patient prognosis and treatment relates on GCT histology and component proportion. In type II GCTs, for example, gGCTs are less aggressive and respond better to chemo and radiotherapy *versus* ngGCTs and therefore require less intensive treatment [1, 10, 11]. The presence of EC alone is a predictor for occult metastasis [2, 8, 10]. Likewise, serum markers alpha-fetoprotein (AFP) and beta-human chorionic gonadotropin (ß-hCG) are associated with YST and CHC, respectively, and elevated levels are linked to worse prognosis [7]. CHC, in particular, is highly aggressive and can metastasize through the blood vessels, bypassing the retroperitoneal lymph nodes [2]. For type I GCTs, it is important to identify and distinguish between TE and YST, as TE are in principle benign but resistant to chemotherapy and thus require surgical removal while YSTs are malignant and often require chemotherapy [10].

Despite overall high survival, GCT patients can suffer from long-term side effects introduced by chemo and/or radiotherapy, such as infertility, secondary malignancies, reduced renal and lung function, and impaired hearing [10]. As such, personalized medicine based on tumor composition and staging is essential to prevent over/undertreatment. There, biomarkers play a crucial role in decision making for optimal clinical GCT patient management. Firstly, immunohistochemistry-based biomarkers are informative for GCT diagnosis and identification of subtypes. Examples include biomarkers for gGCTs (OCT-3/4, POU5F1, SOX-17), EC (OCT-3/4, POU5F1, SOX-2), CHC (ß-hCG, GATA-3), and YST (AFP, FOXA2) [12]. TE are mostly composed of heterogenous cell types and can be generally identified by morphology alone. However, these histomorphological analyses require a tissue biopsy, which is not always possible in a clinical setting. A liquid biopsy (LB) is the sampling of bodily fluids (e.g. blood, bone marrow, and cerebrospinal fluid) that could provide an alternative source for measurement of cancer biomarkers. They are often less invasive to obtain and could mitigate temporal and spatial cancer heterogeneity, hence could provide several advantages over tissue biopsies [13, 14]. Specifically in GCTs, the serum tumor markers AFP, β-hCG, and lactate dehydrogenase (LDH) are proven to be informative and currently considered clinical standards for GCT diagnosis, risk stratification, and follow-up [15]. While AFP and β-hCG are associated with GCT subtypes, LDH is less specific but informative as indicator for disease burden. However, all these markers can also be elevated in conditions unrelated to GCTs [16]. Hence, the lack of specificity and low sensitivity of these serum markers limits utility for histological subtype discrimination [17]. microRNAs of clusters miR-371-373 and miR-302-367 have been shown to be promising alternatives to the classic serum markers [18]. They are detectable in LBs as highly sensitive and specific (elevated in all GCT types except TE) biomarkers informative during all stages of care [19]. However, despite its advantages, these microRNAs are not able to distinguish among different malignant (non-TE) GCT subtypes.

Unlike most other solid cancers in adults, GCTs do not show highly recurrent somatic driver mutations. *KIT, KRAS*, and *NRAS* mutations are the most common (<2 % in Type II ngGCTs of the testis and <20% in gGCTs) [20, 21]. This is explained by the origin of GCTs, through reprogramming of an embryonic germ cell failing to control the latent developmental potential [1]. Epigenetics are thought to play an important role in this reprogramming, GCT onset, and progression [22-24]. DNA methylation is the most studied epigenetic mechanism and has been suggested to be widely involved in GCT onset and pathogenesis [1, 12]. Hence, it provides directions into several GCT directions of study including biomarkers relevant for diagnosis, subtyping, staging, prognosis, therapy response, and epigenetic drugs [22]. Indeed, several methylation-based GCT biomarkers have been proposed, such as *RASSF1A, DPPA3, XIST, CRIPTO, HOXA9, MGMT*, and *SCGB3A1* [25-28]. However, true subtype-specific GCT methylation biomarkers have been lacking so far, most notably because of limited data sets allowing high throughput approaches.

This study provides a pipeline for identification and validation of GCT subtype-specific methylation biomarkers. We performed a meta-analysis on existing and original GCT tissue methylation datasets, based on Illumina 450K and EPIC arrays, including samples from 702 unique GCT patients derived from 15 independent datasets. Using differential methylation analysis tools, we identified common histology-specific biomarkers, of which, as proof of concept, two YST-specific biomarkers were validated in a GCT DNA cohort, and one in an additional cell-free DNA (cfDNA)/LB GCT cohort using methylation-sensitive restriction enzyme (MSRE)-based qPCR.

## Materials and methods

### Meta-analysis study design

The described pipeline for identification of subtype specific GCT methylation-biomarkers is based on a meta-analysis of GCT tissue methylation data generated using Illumina Infinium Beadchip arrays. Published datasets were identified and accessed via the Gene Expression Omnibus (GEO), the National Cancer Institute Genomic Data Commons Data Portal (GDC Data Portal), The DNA Data Bank of Japan (DDBJ), the Functional Genomics Data Collection (ArrayExpress), and on a personal collaboration basis. The following search query was used for identification of GEO datasets that included GCTs: *“(germ cell tumor*^*\**^ *OR germ cell tumour*^*\**^*) AND methylation*^*\**^ *AND (yolk sac tumor*^*\**^ *OR yolk sac tumour*^*\**^ *OR teratoma*^*\**^ *OR choriocarcinoma*^*\**^ *OR embryonal carcinoma*^*\**^ *OR seminoma*^*\**^ *OR germinoma*^*\**^ *OR dysgerminoma*^*\**^*) AND (450K*^*\**^ *OR 450k*^*\**^ *OR EPIC*^*\**^ *OR infinium*^*\**^ *OR illumina*^*\**^ *OR methylation array)”*. Only datasets containing at least one GCT patient-derived primary tumor sample that was profiled by either 450K array or a newer version (EPICv1 and EPICv2) were included. No datasets profiled with EPICv2 were identified. Ultimately, we collected 10 published GCT-related datasets this way. In addition, we provided three original GCT-related datasets. Finally, we included healthy peripheral blood (PB) samples from two additional published datasets. Details per selected dataset are indicated in Table 1.

**Table 1:** overview of included datasets in our meta-analysis. Per dataset, methylation array platforms, the number of included samples, context of the dataset, data source, and reference (if relevant) are provided. Abbreviations: TGCT: testicular germ cell tumor, CNS: central nervous system, GEO: gene expression omnibus, GDC: the National Cancer Institute Genomic Data Commons, DDBJ: national data bank of Japan.

| Dataset ID | Platform | #samples post-screening | Context | Data source | Reference to publication(s) |
| --- | --- | --- | --- | --- | --- |
| 1 | 450K | 213 | Type II TGCTs | Published in GEO (GSE74104) | [26] |
| 2 | EPICv1 | 61 | General GCTs | Original data (GSE303824) | N/A |
| 3 | 450K | 54 | General GCTs | Original data (GSE303825) | N/A |
| 4 | 450K | 143 | Type II TGCTs | Published in GDC Data Portal (TCGA-TGCT) | [20, 29] |
| 5 | 450K | 4 | Pediatric CNS tumors | Published in GEO (GSE215240) | [30] |
| 6 | EPICv1 | 9 | GCT somatic-type malignancy | Published in GEO (GSE219033) | [31] |
| 7 | EPICv1 | 17 | Pediatric GCTs | Original data (GSE303826) | N/A |
| 8 | EPICv1 | 10 | Pediatric CNS tumors | Published in GEO (GSE215240) | [30] |
| 9 | EPICv1 | 51 | Pediatric GCTs | Published in DDBJ (JGAS000204) | [32] |
| 10 | 450K | 145 | Pediatric GCTs | Published in GEO (GSE183798) | [33, 34] |
| 11 | 450K | 92 | Intracranial GCTs | Beta-values published in GEO (GSE70783). Raw data (IDAT) provided on collaborative basis. | [35-37] |
| 12 | 450K | 68 | General GCTs | Published in GEO (GSE58538) | [38] |
| 13 | EPICv1 | 12 | Healthy pediatric blood | Published in ArrayExpress (E-MTAB-13583) | [39] |
| 14 | 450K | 209 | Healthy adult blood | Published in GEO (GSE128235) | [40] |
| 15 | EPICv1 | 4 | GCT somatic-type malignancy | Published in GEO (GSE281147) | [41] |
|  | Total | 1092 |  |  |  |

For all identified datasets, human GCT samples were only included for *in silico* analysis when they were derived from a primary GCT location (being non-metastasized) and had a unique sentrix ID (avoiding duplicate samples). In addition, datasets 1, 2, 3, 4, 12 contained patient overlap despite having unique sentrix IDs, as samples within these datasets came from the same institute (Erasmus MC, Rotterdam, the Netherlands). If this was the case, the following order was maintained for final sample selection: 1>4>12>3>2. As such, only samples from true unique patients were retained. The only exceptions were if a mixed tumor was dissected into multiple separate histologic components that were profiled separately (then all individual components were included as samples), and patients from which both tumor tissue and benign neighboring testis (BNT) was profiled (only relevant for dataset 1). We also included seven GCT cell lines (CLs): TCam-2 (representing SE), NT-2, 2102Ep, and NCCIT (all three EC), NOY-1 (YST), JEG-3 and JAR (both CHC). For all cell lines (except TCam-2, parental only), parental and cisplatin sensitive clones were included to screen for potential cisplatin biomarkers. Finally, any sample (tissue, CL, and PB) was only included for analysis when passing QC metrics (see DNA methylation integration pipeline in the M&M section for details). Following this selection procedure, our final dataset included 822 tissue samples (713 GCT and 109 BNT) derived from 702 unique GCT patients, 49 CL samples, and 221 PB samples.

### Patient samples

For patient-derived tumor tissue samples (from datasets 2 & 3 of the meta-analysis cohort, and the entire MSRE qPCR validation cohort): the use of leftover tissue material for scientific purposes was approved by the Medical Ethical Committee (MEC) of the Erasmus MC Rotterdam (Rotterdam, the Netherlands), permission 02.981. For samples from dataset 7 of the meta-analysis cohort: use was approved by the Biobank and Data Access Committee (BDAC) of the Princess Máxima Center (Utrecht, the Netherlands), permission 2020.181 for intracranial and 2020.182 for extracranial GCT tumors. Tumor samples were either formalin-fixed and paraffin-embedded (FFPE) or freshly frozen, and subsequently stored in liquid nitrogen. For patient-derived serum samples, blood was collected before treatment between 1996 and 2018 in several Dutch hospitals, and serum was stored at -80°C (also permission 02.981 by the Erasmus MC Rotterdam). Detailed metadata for all three cohorts are provided in Table S1 (for meta-analysis cohort), S2 (MSRE qPCR validation cohort), and S3 (LB cohort).

### DNA isolation and methylation arrays

DNA isolation, bisulfite conversion, and methylation profiling of patient and CL-derived tumor DNA using Illumina’s 450K and EPICv1 arrays for generating original datasets of the meta-analysis cohort (datasets 2, 3, and 7) was done as previously described [25, 38, 42, 43]. Patient-derived tumor DNA isolation for the MSRE qPCR validation cohort was done the same way.

For patient serum samples (used in the LB cohort), cfDNA isolation was performed using the Maxwell RSC ccfDNA LV Plasma Kit (AS1840, Promega) according to manufacturer’s protocol. Following cfDNA isolation, a double size-selection step using AMPure XP Beads (Beckman Coulter) was performed to remove any genomic DNA. For conditioned medium of GCT CLs, 2 mL medium was taken, centrifuged at 3000xg, and DNA isolation was performed using the Maxwell RSC ccfDNA Plasma Kit (AS1480, hence not LV kit as with serum) according to manufacturer’s protocol.

### Cell line culture

Eight different GCT cell lines were cultured for this study: TCam-2, NT-2, 2102Ep, NCCIT, GCT-44, NOY-1, JEG-3, JAR. Culture conditions were as previously described [37].

### Meta-analysis

#### Pre-analysis pipeline

R (v4.4.2) was used for all data (pre)processing and analysis. Per dataset, raw IDAT files were read and converted to an RGChannelset using read.metharray.exp function of the minfi package (v1.52.1) [44]. Outlier samples and probes were identified using the bscon and pfilter functions (QC checks 1) of the wateRmelon package (v2.12.0) and excluded [45]. RGChannelset formats were preprocessed using the preprocessFunnorm function and converted to a GenomicRatioSet using minfi [46]. Beta-values were calculated using the getBeta function and merged together from the 15 individual datasets. Probes not shared among the 450K and EPICv1 arrays, SNP probes, cross-reactive probes, and sex chromosome-specific probes were removed using the rmSNPandCH function of the DMRcate package (v3.2.1) (QC check 2) [47]. Missing values were imputed using the methyLImp2 function of the same package (v1.2.0). Then, beta-values were corrected for signaling specifically introduced by lymphocytes (see details below). Finally, beta-values were converted to M-values where beta-values of 0 and 1 were replaced with 1e-6 and 1-1e-6 to prevent division by zero or infinity, respectively. Our final dataset included n=1,092 samples with n=404,246 probes annotated to unique CpG sites within the human genome. A graphical overview of the pre-analysis pipeline is included in Figure 1.

**Figure 1:**
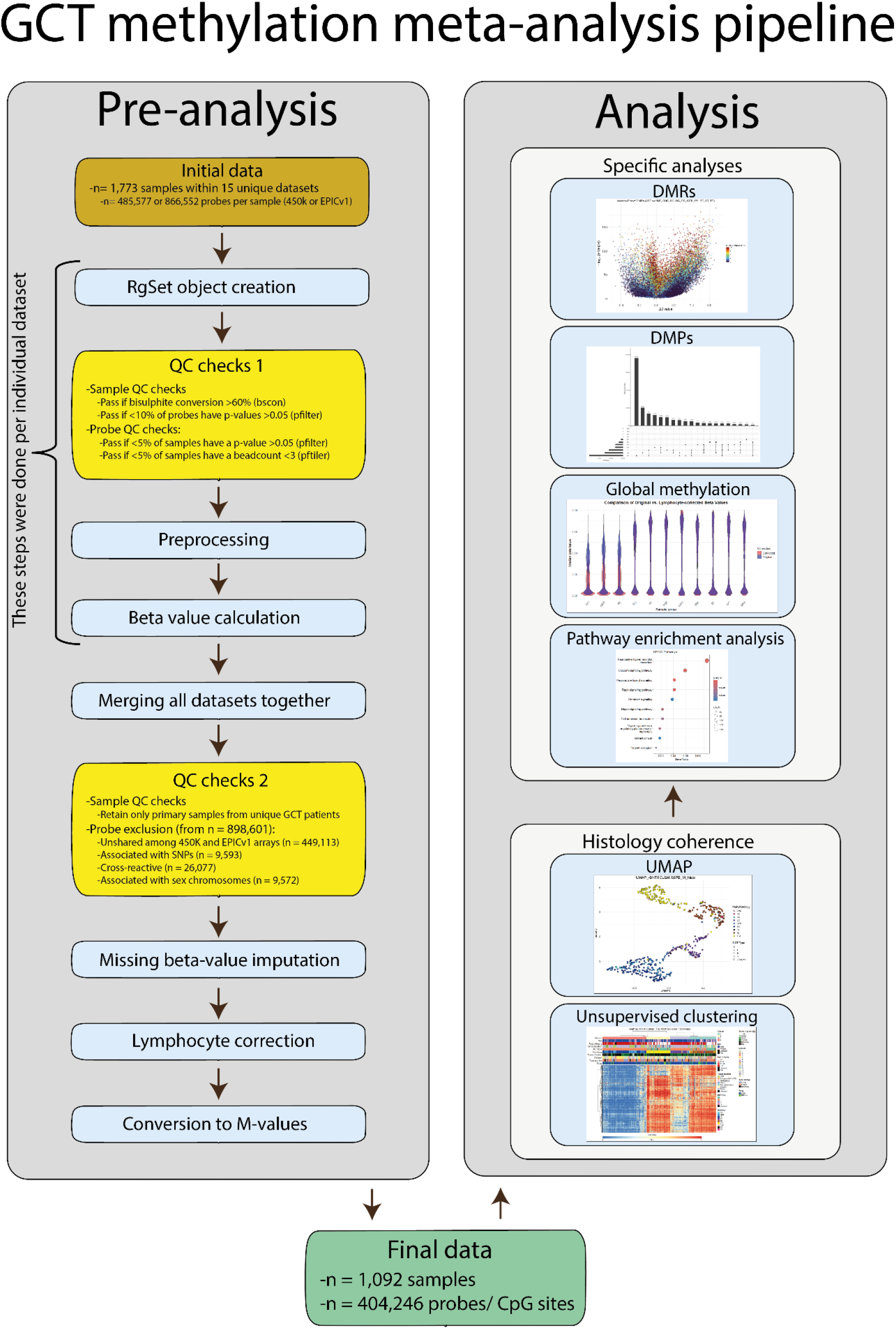
graphical overview of the full meta-analysis pipeline for in silico identification of GCT subtype-specific biomarkers. Key steps for data pre-analysis and QC are indicated in the left column. Final data was first checked for histology coherence using UMAP and UHC followed by more specific analyses (DMRs, DMPs, global methylation profiling and pathway enrichment analysis) (right column). Abbreviations; QC: quality control, UMAP: Uniform Manifold Approximation and Projection, DMR: differentially methylation region, DMP: differentially methylated position.

#### Lymphocyte correction and global methylation profiling

We used a correction method developed and applied by Killian *et al* [26]. This includes a lymphocyte cell-specific hypermethylation module (36 selected probes based on lymphocyte (n=18) and non-lymphocyte (n=87) reference data) to quantify the degree of lymphocyte infiltration per sample (referred to as the lymphoid index (LI)). A sample’s LI was defined by taking the average beta-value of those specific 36 probes, bounded between 0-1. Next, a two-component model assuming that an observed beta-value consists of a lymphocyte and non-lymphocyte signal was applied for lymphocyte correction: (*β*_1_=LI∗*β*_LY_+(1-LI)∗*β*_2_). There, per probe of a specific sample: *β*_1_=the observed beta-value before correction, *β*_2_=the corrected beta-value/non-lymphocyte signal, *β*_LY_=the average beta-value for the 18 reference lymphocyte samples/lymphocyte signal, LI=that sample’s LI index (the same per sample regardless of the probe)). This formula is rearranged into the following: *β*_2_=(*β*_1_−LI∗*β*_LY_)/(1−LI). This correction method was applied to all tissue samples (hence not PB and CL samples). For global methylation, median beta-values per CpG site were calculated among all samples, per histology group. Then, the mean of these medians was calculated to obtain a single global methylation number, per histology group. Violin plots were generated using the ggplot2 package (v3.5.2).

#### PCA and UMAP

Principal component analysis (PCA) was first performed on M-values, using the base R prcomp function without scaling. PCs explaining 0.90 of the cumulative variation were included for Uniform Manifold Approximation and Projection (UMAP), which was performed using the umap function of the umap package (v0.2.10.0) using a fixed seed (123456789) [48]. Plots were generated using ggplot2.

#### Unsupervised clustering

Out of the total 404,246 CpG sites, the top 1000 displaying the highest median absolute deviation were identified. Unsupervised hierarchical clustering (UHC) was performed on beta-values of this top 1000 using the ConsensusClusterPlus function (reps=10,000, clusterAlg=hc, distance=euclidean, innerLinkage=ward.D2, finalLinkage=ward.D2, seed=123456789) of the ConsensusClusterPlus package (v1.70.0) [49]. The optimal amount of clusters was chosen based on visual inspection of the Consensus Cumulative Distribution Function (CDF) Delta Area Plot (highest k that still results in an appreciable decrease in relative change in area under the CDF plot). Dendrograms were generated for samples (columns) based on the UHC results, and for CpG sites (rows) using the hclust function (baseR, method=ward.D2). The final heatmap was plotted using the Heatmap function of the ComplexHeatmap package (v2.22.0).

#### Differentially methylated position analysis

Differentially methylated positions (DMPs) represent single genomic CpG sites with a statistically significant difference in methylation between two groups. Per GCT histology, all samples were statistically tested (F-test using beta-values) against all other samples in the total cohort (excluding CLs and MIX samples) using the dmpFinder function of the minfi package (type=categorical). P-values were corrected for multiple testing according to the Benjamini-Hochberg method. Results were expressed as volcano plots and upset plots (UpSetR package v1.4.0) with data points representing unique CpG sites based on significance (false detection rate (FDR)-corrected p-values) and mean differences in beta-values (Δbeta).

#### Differentially methylated region analysis

Differentially methylated regions (DMRs) represent genomic regions containing at least 2 CpG sites with a statistically significant degree of methylation between two groups. Per GCT histology, all samples were tested against all other samples in the total cohort (excluding CLs and MIX samples) for DMR analysis. This was done using the cpg.annotate, dmrcate, and extractRanges functions (datatype=array, coef=2, arraytype=450K, what=M, analysis.type=differential, lambda=1000, C=2, genome=hg19) of the DMRcate package. Plots were generated using the DMR.plot function [47]. Results were expressed as volcano plots with data points representing individual significant DMRs based on HMFDR (harmonic mean of the FDR-corrected individual CpG p-values that make up a DMR), Δbeta, and length (number of CpGs within the DMR).

### Pathway enrichment analysis

Following group-specific DMR analyses, DMRs with a Δß-value >0.2 and <-0.2 were taken (to filter out the most biologically relevant ones) and involved genes and their genomic feature annotations were extracted using the CHIPseeker package (v1.42.1) with the TxDb.Hsapiens.UCSC.hg19.knownGene transcript database and org.Hs.eg.db annotation. Gene symbols were extracted and mapped to Entrez IDs. The background gene universe consisted of all genes present in our final Illumina probe/CpG set. KEGG pathway enrichment analysis was performed using the enrichKEGG function of the clusterProfiler package (v4.14.6) with a p < 0.01 threshold and Benjamini-Hochberg correction. Pathway enrichment using the Panther metabolic and cell signaling pathway database (Panther 2016) was done in a web browser using enrichR (https://maayanlab.cloud/Enrichr/).

### MSRE-based qPCR assay development for detection of methylated *APC* and *DPP7*

MSRE-based qPCR was chosen for targeted detection of methylated DNA as it is fast, cheap, and simple, and requires minimum DNA input *versus* alternative (bisulfite conversion-based) methods [50, 51]. Primers were designed (Geneious Prime software) around specific *APC* and *DPP7* gene regions that were identified by the DMR analysis pipeline (see Table S4 for sequences). Target amplicons should cover at least 1 CpG site included as probe within our meta-analysis dataset. Primer pair amplification efficacy and specificity were tested using endpoint PCR followed by gel electrophoresis and qPCR followed by melting curve analysis. The methods for MSRE qPCR assays described below refer to the final optimized protocol. Per experiment, four PCR mastermixes were prepared (*APC/DPP7*, +/-restriction enzymes). Master mixes contained *APC* or *DPP7* primer pairs (integrated DNA technologies), SYBR green dye (Thermofisher), Milli-Q, and either a mix of MSREs (AciI + HhaI for *APC*, and HpaII + HhaI for *DPP7*) or Diluent A (all from New England Biolabs). 20ng total DNA was used as input per condition. The qPCR protocol contained the following steps: 15 minutes at 37°C (enzyme activation and digestion of unmethylated target DNA) -> 10 minutes at 95°C (enzyme deactivation) -> 40 cycles of 15 seconds at 95°C + 1 minute at 60°C (DNA amplification). Per sample, the ΔCt value was calculated (Ct of condition with enzyme – Ct of condition without enzyme). A standard curve containing different ratios of GCT cell lines (GCT-44 and JAR as positive methylation controls for *APC* and *DPP7* respectively, and NOY-1 as negative methylation controls for both targets) was used to convert ΔCt to a methylation % per test sample. Graphs were made using GraphPad Prism 10.

## Results

### Cohort description

In total, 702 unique GCT patients (Table 2) were included in our meta-analysis cohort. They represent patients from a wide age range (0-82 years, median=19), broadly divided into prepubertal (0-7 years, 20%), pubertal (8-18 years, 26%, and post pubertal (>18 years, 50%) categories. Most patients were male (75%), and most patients had a gonadal primary tumor (70%). Pure primary GCTs were divided into 10 histology groups: CHC, dermoid cyst (DC), DG, EC, GER, GCT with mixed components (MIX), SE, spermatocytic tumor (ST), TE, and YST. Outcome data was not available for many datasets (included in Table S1 if available) and hence has not been included as parameter.

**Table 2:**
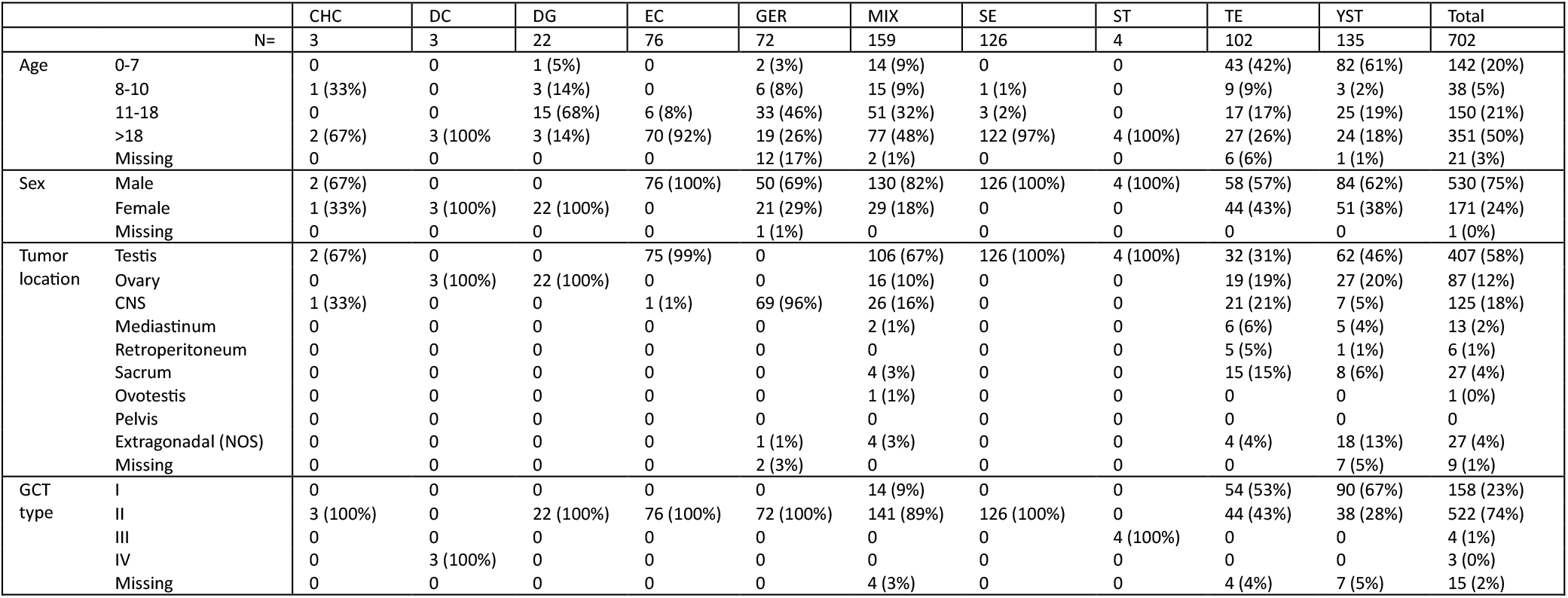
clinical and histological characteristics of unique patients within our meta-analysis cohort by histologic subtype. Abbreviations: CHC: choriocarcinoma, DC: dermoid cyst, DG: dysgerminoma, EC: embryonal carcinoma, GER: germinoma, MIX: mixed germ cell tumor histologies, SE: seminoma, ST: spermatocytic tumor, TE: teratoma, YST: yolk sac tumor.

### GCT methylation clustering

We utilized UHC to investigate histology coherence and identify clusters with shared global methylation profiles within our meta-analysis cohort. When analyzing only histologically pure samples, four distinct clusters were identified (Figure 2, Table S5). Clusters 1, 2, and 3 included most TEs (99%, 102/103), YSTs (83%, 112/135), and ECs (91% 71/78), respectively, each with distinct methylation patterns. Cluster 4 included the majority of gGCTs (DG: 86% (19/22), GER: 84% (64/76), SE: 98% (125/128)), exhibiting a hypomethylation profile. All CHC and DC samples (both 3/3) clustered within the TE cluster, while all (4/4) ST samples clustered within the EC cluster. Apart from histology, other patient (sex, age) and tumor (anatomical location, GCT type (for YST and TE only), tumor fraction, and archive type) parameters were not associated with any specific clusters. Also, absence of clusters associated with dataset ID or array type argue against the presence of any batch effects.

**Figure 2:**
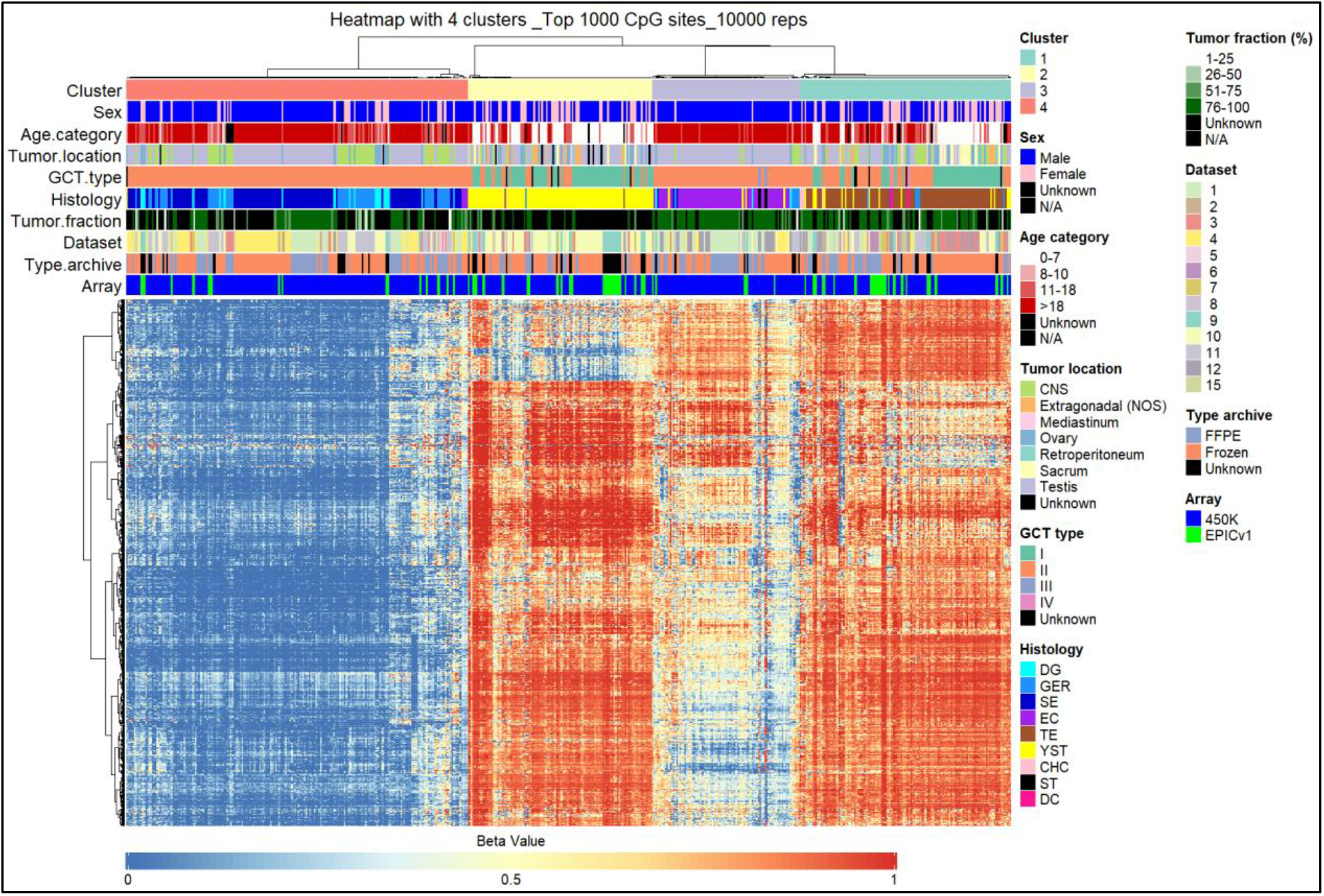
GCTs with the same pure histology generally cluster together regardless of other parameters. Methylation-based UHC of GCT samples with pure histology was performed on beta-values, using the top 1000 CpG sites with highest median absolute variation and 10,000 repetitions. Heatmap columns represent unique samples, rows represent unique CpG sites. Core data represents beta-values ranging from 0 (fully demethylated) to 1 (fully methylated) based on a color gradient scale. Annotations with several clinical and histological parameters are provided on top with corresponding legends provided on the right. The amount of clusters was chosen based on visual inspection of the corresponding Delta area plot (Figure S8). Abbreviations: CHC: choriocarcinoma, DC: dermoid cyst, DG: dysgerminoma, EC: embryonal carcinoma, GER: germinoma, SE: seminoma, ST: spermatocytic tumor, TE: teratoma, YST: yolk sac tumor

Subsequently, we included mixed GCTs (MIX), benign neighboring testis (BNT), and GCT cell lines (CL) in the UHC pipeline. Inclusion of these groups preserved the four primary clusters corresponding to gGCTs, ECs, TEs, and YSTs across all conditions. MIX samples were distributed among all four clusters (Figure S1, Table S6). Of all MIX samples, 19% (31/161) had a major histological component (defined as ≥75% contribution to total histology). For those, 100% (12/12) of MIX_EC, 75% (6/8) of MIX_TE, 75% (3/4) of MIX_SE, and 57% (4/7) of MIX_YST cases clustered with their respective pure histology counterparts. Most BNT samples were distributed across the TE (42%, 46/109) and EC (55%, 60/109) clusters (Figure S2, Table S7). Johnson score (JS) is a scoring system for spermatogenesis ranging from 0 (absence of germ and being Sertoli cell only) to 10 (complete spermatogenesis with intact tubules) [52, 53]. JS data (available for dataset 1 only) was categorized into zero/low, and mid/high groups [26]. We observed a significant association (p=1.1e^-12^, Fisher’s exact test) between BNT clustering and JS where BNT samples with mid/high JS had a tendency to cluster with EC samples while BNT samples with zero/low JS generally clustered with TE samples. For GCT CLs, TCam-2 clustered with EC tumors while all other CLs (2102Ep, NCCIT, NT2, JEG-3, JAR, and NOY-1) clustered with TE histology (Figure S3).

To further assess sample similarity based on methylation profiles, we applied UMAP. In contrast to UHC, UMAP preserves local data structure for assessing sample heterogeneity within groups [48]. Overall, UMAP results were consistent with the UHC, demonstrating robustness of histology clustering. For pure GCT samples, UMAP revealed clear segregation of the majority of gGCTs, ECs, TEs, and YSTs (Figure 3). No clear separation was observed between type I and II TEs or YSTs. For EC and TE samples, subgroups distinguishing FFPE from frozen samples were identified (Figure S4A). Inclusion of MIX samples roughly maintained the YST and gGCT groups but increased distribution of TE and EC samples, with MIX samples appearing within and between pure GCT groups (Figure S5). Inclusion of BNT samples resulted in a new BNT group together with a subset of TE samples while the rest of the TEs, YST, gGCT and EC groups were maintained (Figure S6). Finally, inclusion of GCT CLs resulted in CL localization into three separate groups, entirely distinct from the GCT samples (Figure S7).

**Figure 3:**
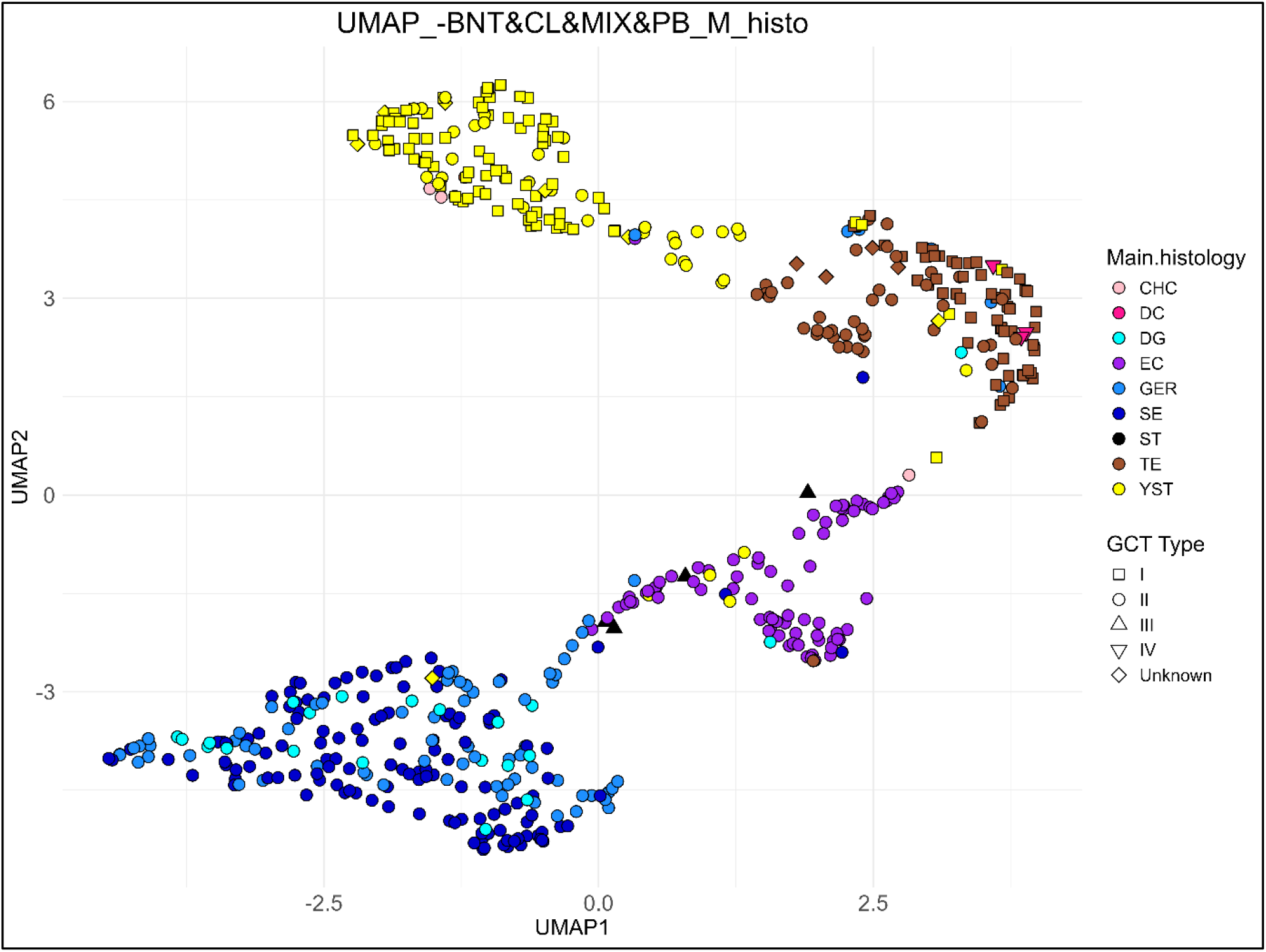
GCTs with the same pure histology generally group together regardless of other parameters. UMAP of GCT samples with pure histology was performed on M-values. Samples are color-coded by their histology. Data point shapes refer to GCT type (relevant only for YST and TE samples). Abbreviations: CHC: choriocarcinoma, DC: dermoid cyst, DG: dysgerminoma, EC: embryonal carcinoma, GER: germinoma, SE: seminoma, ST: spermatocytic tumor, TE: teratoma, YST: yolk sac tumor.

Taken together, the results from hierarchical clustering and UMAP indicate that histology is the primary determinant of a GCT’s methylation profile, independent of a patient’s sex, age, and tumor location.

### Lymphocyte-compensated GCT histology-specific global methylation profiling

Patient derived tumor samples generally consist of not only tumor cells, but also other cell types such as immune and stromal cells representative of the tumor microenvironment [54]. Consequently, mixtures of such cell types can affect the methylation profiling readout (beta-values) [55]. Specifically in the context of GCTs, particularly gGCTs often contain various amounts of infiltrative lymphocytes. [20, 26, 56-60]. As gGCTs are generally hypomethylated, their profile could be (partly) masked if lymphocyte infiltration is high [20, 26, 32, 34, 35, 38].

We calculated the lymphoid index (LI) to estimate the degree of lymphocyte infiltration of the samples in our meta-analysis cohort [26, 61]. We observed a LI of 0.34, 0.44, 0.40 for gGCTs (DG, GER, and SE, respectively) and 0.61, 0.34, 0.23, 0.25 for ngGCTs (CHC, EC, TE, and YST) (Figure S9). In line with previous studies, this illustrates that both gGCTs and ngGCTs can exhibit lymphocyte infiltration, and potential effects need to be considered during GCT methylation analyses [20, 26]. Therefore, we performed a correction to mitigate bias in methylation profiling introduced by lymphocyte infiltration (see methods for details) as described before [26, 61]. Lymphocyte correction resulted in a decrease of median beta-values for gGCTs (-0.21, -0.25, -0.21 for DG, GER, SE) while the median beta-values for ngGCTs remained similar (-0.02, -0.06, -0.03, -0.02 for CHC, EC, TE, YST) (Figure S10). All GCT histologies displayed a hypomethylation peak (defined here as a high probe density with beta-values ≤0.25) that was typically more dense in gGCTs *versus* ngGCTs. Additionally, ngGCTs also displayed a hypermethylation peak (beta-value: ≥ 0.75), which was absent in gGCTs. Concurrently, the lymphocyte-corrected global methylation profile for gGCTs was significantly lower *versus* the other GCTs (EC, TE, YST, CHC, MIX, ST, DC) (0.17 +/-0.02 *versus* 0.44 +/-0.04, p-value < 0.01, Mann-Whitney U test) (Figure S10). In conclusion, our analysis further supports the notion that gGCTs display a typical globally hypomethylated signature *versus* ngGCTs.

### Identification of GCT histology-specific methylation profiles

As GCT histology seems the main driver for a sample’s methylation pattern, we proceeded with identification of histology-specific methylation profiles. We identified specific significantly hyper/hypomethylated DMPs/CpG sites (expressed as probes having a FDR-corrected p-value <0.05 and a Δbeta >0.2 or <-0.2), and investigated common sample groups sharing significant probes (Figure S11). The PB group contains the most exclusively hypermethylated probes (95,255, 24% of all probes) followed by YST (Figure S12). In contrast, most hypomethylated probes are shared by the gGCTs with 151,711 CpG sites (37%) being exclusively hypomethylated in this group, and 175,896 CpG sites (44%) being exclusively hypomethylated in one or more gGCT types (Figure S12).

While DMPs are informative, differentially methylated regions (DMRs) are more biologically relevant as they reflect changes in methylation across genomic regions. We identified numerous hyper- and hypomethylated DMRs per histology group (Table S8, Figure 4A and Figure S13). gGCTs were treated as a single group (DG/GER/SE) based on the overlap in individually hypomethylated CpG sites (Figure S12). TE, YST, EC, and CHC displayed a higher number (9,260-31,391) of hypermethylated DMRs compared to ST, DG/GER/SE, and DC (10-2,168). gGCTs, unsurprisingly, displayed the highest number (44,926) of hypomethylated DMRs, with other groups displaying a lower number (7-10,128). A detailed overview of GCT subtype-specific DMRs per GCT subtype (and group comparisons) is provided in Table S9-17. DMRs can link to potential biomarkers. An example of this *in silico* identification pipeline is provided for YSTs in Figure 4.

**Figure 4:**
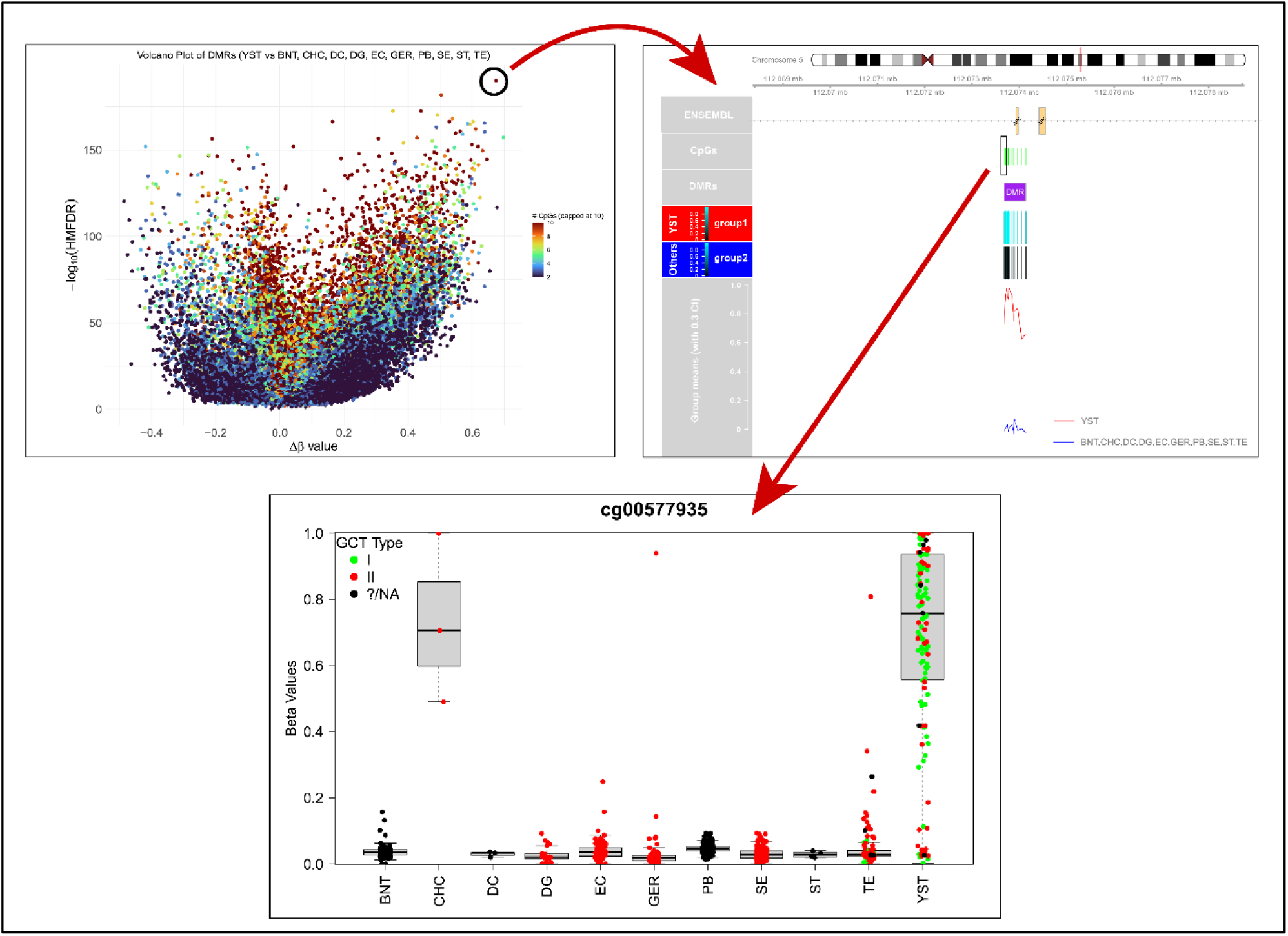
example of in silico GCT subtype-specific biomarker identification workflow. All YST samples were tested versus all other samples in the meta-analysis cohort (excluding MIX and CLs). Datapoints represent unique DMRs with positive Δß-values representing hypermethylated YST-specific DMRs and negative values hypomethylated YST DMRs (A). For an example DMR, the mean Δß-values for all collectively included probes are plotted for both groups (B). Also, relevant genomic coordinates are provided. For an example probe/CpG site within the DMR, the beta-values are plotted for all individual samples per group (C). Abbreviations: BNT: benign neighboring testis, CHC: choriocarcinoma, DC: dermoid cyst, DG: dysgerminoma, EC: embryonal carcinoma, GER: germinoma, PB: peripheral blood, SE: seminoma, ST: spermatocytic tumor, TE: teratoma, YST: yolk sac tumor.

Previously, we identified *DPPA3* hypomethylation and *RASSF1A* hypermethylation as collective GCT markers [25, 26]. These markers were confirmed in our meta-analysis (Table S17). For *DPPA3*, three DMRs were identified of which one was ranked 24 out of 44,698 when sorting collective GCT DMRs by hypomethylation (from low to high mean Δß). Similarly, for *RASSF1A*, three DMRs were identified of which one ranked 366 out of 44,698 when sorting DMRs by hypermethylation (from high to low Δß). Example probes annotating to *DPPA3* and *RASSF1A* are indicated in Figure S14.

Our meta-analysis cohort includes several parental and cisplatin-resistant GCT CLs. We aimed to also identify DMRs between collective parental and cisplatin resistant GCT CLs as potential markers for cisplatin resistance, as this is the most important predictor for poor outcome in GCT patients, but did not identify any (data not shown) [1].

### Pathway enrichment analysis

DMRs can be mapped to genomic coordinates to identify corresponding gene(s) and their genomic feature annotations. Then, enrichment analysis on the resulting gene list allows for identification of biologically relevant pathways. Between 1,621 and 12,074 differentially methylated genes were identified for GCT subtypes (excluding DC where five were identified) (Table S18). For all subtypes, most genes (26-47%, again excluding DC) were featured on promotor regions (<=1kb upstream from translational start site) (Table S18). Also, most genes were shared among multiple GCT subtypes except for the gGCTs for which 29.8% (3,602/12,074) were not shared with any other group (Table S18 and Figure S15). KEGG enrichment of these gene sets, per histology, revealed several pathways for CHC, gGCTs, and YSTs of which many were conserved, most likely due to gene overlap (Figure S16A-C). As such, we compared mutually exclusive genes for collective gGCTs and ngGCTs revealing only two gGCT but numerous ngGCT-specific KEGG pathways (Figure S16D-E). While KEGG enrichment on mutually exclusive genes for individual ngGCT subtypes did not result in any enriched pathways, enrichment using the Panther 2016 database revealed the WNT and Cadherin signaling pathways as YST-specific (Figure S16F). Similar to KEGG, no significant CHC, EC, an TE-specific pathways were identified by Panther 2016. Gene lists per histologic analysis are provided in Table S19.

### MSRE qPCR assay development for quantification of methylated *APC* and *DPP7* as YST-specific biomarkers

Out of all GCT subtypes, YST showed the most significant hypermethylated DMRs (based on Δbeta, log10(HMFDR), and cohort size (Table S8)). Hence, we focused on this subtype, and identified the two most significant (one with highest Δbeta, and one with highest log10(HMFDR)) YST DMRs, which contained 15 and 5 CpG-annotated probes within or surrounding the promoter regions of *APC* and *DPP7* genes. We developed MSRE qPCR assays for specific detection of *APC* and *DPP7* methylation, serving as proof of concept validation of methylation-based biomarker identification by our *in silico* pipeline. MSRE qPCR assays involve a restriction enzyme reaction mix resulting in digestion of only unmethylated DNA leaving methylated DNA intact. Following digestion and subsequent enzyme inactivation by heat, only intact (and thus methylated) DNA target copies are amplified using qPCR. The resulting ΔCt (which is calculated using an undigested control condition) is a quantitative measure for degree of digestion that is correlated to a methylation % using a standard curve.

Several conditions were optimized related to target amplicon (primer specificity), restriction enzymes (combinations, units of input, incubation time, inactivation conditions), DNA input, and limit of quantification (LOQ). Briefly, primer pairs were designed for different amplicons within the *APC* and *DPP7* DMRs, with the best pair per target selected based on gel electrophoresis and melting curve analyses (Figure S17). Assays were optimized using GCT CLs TCam-2 (hypomethylated) and JEG-3 (hypermethylated) (Figure S18). Optimal digestion (highest TCam-2 without JEG-3) was achieved using 1 unit (U) of AciI + 1U HhaI *APC*, and 1U HpaII + 1U HhaI for *DPP7*, both with 20 ng total DNA input (hence 0.1 total enzyme U/ng DNA), incubated at 37°C for 15 minutes (Figure 5A-B, Figure S19-20). For *APC*, GCT-44 and NOY-1 GCT CLs were selected as methylated and unmethylated controls based on ΔCt, being non-inferior to commercial controls (Figure 5C). For *DPP7*, JAR and NOY-1 were selected. These controls were used to generate standard curves (Figure 5D). Interestingly, similar *APC* and *DPP7* methylation levels were observed in conditioned medium of GCT-44 cells as in GCT-44 cells (not tested for other GCT CLs) (Figure S21). A fully elaborated overview of assay optimization results are provided in Supplemental data 1.

**Figure 5:**
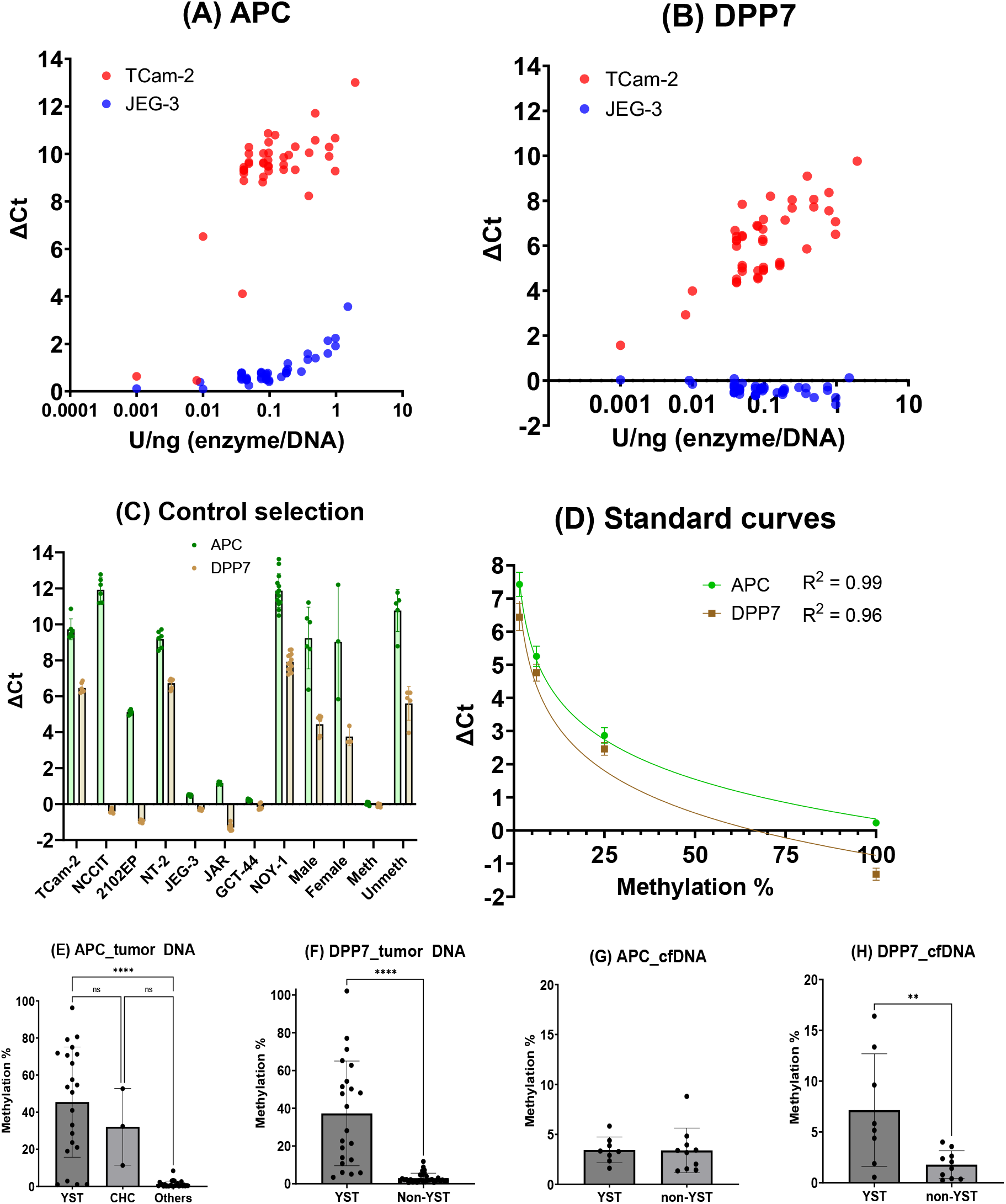
MSRE-qPCR assay development and validation. The effect of MSRE unit (U) input/ng DNA input (x-axis) on APC (A) and DPP7 (B) DNA digestion efficacy is shown for TCam-2 and JEG-3 GCT cell lines. Using an optimized input of 0.1 U/ng, different GCT CLs, healthy control DNA (Male & Female) and commercially available methylated (Meth) and unmethylated (Unmeth) DNA were tested (C). GCT-44/NOY-1 (for APC) and JAR/NOY-1 (for DPP7) were chosen as methylated and unmethylated controls, and used to make standard curves correlating observed ΔCt to a calculated methylation % (D). Assays were validated in tumor DNA (CHC, DG, EC, MTE, SE, YST) and healthy PB for APC (E) and DPP7 (F). Methylation % for individual groups and metadata for E/F is shown in Figure S23 and Table S2. APC (G) and DPP7 (H) methylation was tested in cfDNA of GCT tumor patients and healthy controls. Methylation % for individual groups and metadata for G/H is shown in Figure S25. Metadata for G/H is shown in Table S3. Abbreviations: cfDNA: cell-free DNA, CHC: choriocarcinoma, DG: dysgerminoma, EC: embryonal carcinoma, PB: peripheral blood, SE: seminoma, MTE: mature teratoma, YST: yolk sac tumor.

We proceeded with *APC* and *DPP7* assay validation in tumor DNA from the MSRE qPCR validation cohort. This cohort consists of 53 GCT patients with CHC, DG, EC, TE, SE, or YST histology, as well as PB from six healthy subjects as controls. For *APC*, we observed higher methylation in the YST and CHC groups *versus* all other samples (Figure 5E). This difference was only significant for the YST group versus the other group (Kruskal-Wallis test + Dunn’s multiple comparisons test: p < 0.0001 for YST *versus* others and p=0.0547 for CHC *versus* others), likely due to the small sample size in the CHC group. For *DPP7*, we observed a significantly higher methylation % in the YST group *versus* all other (including CHC) samples (Mann-Whitney test, p < 0.0001) (Figure 5F). For both targets, the results concurred well with the expectation (including the *APC* methylation signal in CHC) based on our meta-analysis (see example probes for *APC* and *DPP7* amplified regions, Figure S22). The methylation % per individual sample group for both targets is indicated in Figure S23.

Finally, we aimed to validate our markers in serum-derived cfDNA from the LB cohort. This cohort consists of 18 GCT patients with CHC, EC, SE, YST, or mixed histology, as well as cfDNA from two subjects without malignancy as additional controls. The tumor DNA-optimized assay requires 80ng of DNA material per experiment, which is unfeasibly high for repeated LB experiments as cfDNA yield is generally low (we had 30–156ng cfDNA available for this cohort). Therefore, we first performed an LOQ experiment. For *APC*, an input of 2ng with 0.2U (thus still providing 0.1 U/ng ratio) was the limit (Figure S24). For *DPP7*, 2ng with 2U (thus 1 U/ng) was the limit. Testing our patient-derived LB samples using this approach revealed a significantly higher *DPP7* methylation % in subjects with a YST *versus* all subjects without a YST component (Figure 5G). For *APC*, no differences were observed (Figure 5H). The methylation % per individual sample group for both targets is indicated in Figure S25.

## Discussion

In this study, we present a pipeline for *in silico* identification and *in vitro* validation of methylation-based biomarkers for subtype-specific human GCTs, based on integration of 15 independent datasets. UHC and UMAP revealed that GCT histology is most strongly associated with the observed methylation profiles. However, not all samples of the same histology clustered together. This was predominantly the case for YST samples where 17% (23/135) clustered elsewhere (mostly either with TEs or ECs). Samples that clustered differently based on UHC generally also grouped differently based on UMAP (Figure S4B). Potential causes include tumor heterogeneity, especially differences in tumor purity, contamination with non-tumor cell types such as immune or stromal cells, or misclassification based on limited or non-representative diagnostic material. As GCTs often occur as mixed histology types, especially in the case of ngGCTs, some samples classified as pure may still contain mixed components [1]. Although most utilized datasets clearly described methods for histology classification, variations in criteria or pathological review may explain some of the observed clustering discrepancies. Also, GCTs can have trajectories of different histologies. As example, EC can differentiate into all three germ layers (TE) and extra-embryonic tissues (YST and CHC), while gGCTs can reprogram into EC or other ngGCTs [1, 62]. Similarly, TEs can progress into YSTs (particularly in type I GCTs) or somatic-type malignancy [41, 63]. UMAP revealed an interesting pattern where a subgroup of gGCT and EC samples grouped closely together and separate from their main groups (Figure 3). Similar patterns were observed between some EC and TE, and YST and TE samples. This aberrant pattern might reflect the developmental potential of GCTs where some samples exhibit differentiating behavior. Several datasets (4, 5, 8, 9, and 10) provided paired RNA expressions, which could help determine whether differently clustered samples of the same histology represent true biological differences or outliers introduced by misclassification or non-pure tumor tissues. Finally, we included different GCT CLs in our clustering analyses of which none reflected the methylation profiles of their respective *in vivo* GCT counterparts (EC lines were still OCT3/4 positive, data not shown), warranting caution when using these cell lines as model systems in GCT methylation studies.

We identified numerous histology-specific DMRs with biomarker potential (Table S9-17). YST-specific hypermethylated DMRs were most significant, leading to selection of *APC* and *DPP7* for proof of concept validation. In tumor DNA, significant *APC* methylation was observed in YST and CHC samples, but not in any of the other groups. This concurs with expectations following our meta-analysis. Significant *DPP7* methylation was observed exclusively in YST DNA. In contrast, our meta-analysis showed *DPP7* methylation in 1/3 CHC samples. The single *DPP7*-methylated CHC samples was not included in the tumor DNA validation cohort as it was derived from an external dataset. More CHC samples are needed to confidently assess whether *DPP7* methylation is truly YST-specific or whether it can also occurs in CHC. However, this can be challenging as pure CHC is extremely rare. Following validation in tumor DNA, we investigated marker performance in cfDNA to explore relevance for LBs. We measured significant *DPP7* methylation in cfDNA of GCT patients with a YST-component, although the difference was less pronounced compared to tumor DNA. Tumor fraction in cfDNA is inherently lower than in tumor DNA as cfDNA is derived from the circulation and hence contains abundant DNA from non-tumor origin [13]. *DPP7* methylation varied across YST-component cfDNA samples, possibly reflecting differences in YST-size, tumor stage, or metastatic status. Future studies with bigger cohorts could correlate *DPP7* methylation levels to these parameters to evaluate clinical utility for different applications including diagnosis, prognosis, treatment response, follow-up, and relapse monitoring. There, *DPP7* methylation should be paired with classical serum markers (AFP) and miRNA-371a-3p analyses to compare performance for each application. *DPP7* methylation was similar in GCT patients without a YST-component and subjects without malignancy indicating presence of background methylated *DPP7* in blood. Review using the Epigenome-Wide Association Study (EWAS) open access platform revealed methylation of probe cg13575271 (being covered by the *DPP7* amplicon) in healthy adipose tissue, which could be a source of methylated *DPP7* in subjects without malignancy (Figure S26) [64]. In contrast to *DPP7, APC* methylation was not elevated in the cfDNA of YST-patients. This indicates that methylation in tumor DNA does not guarantee detectability in cfDNA. cfDNA release (necrosis, apoptosis, and active secretion) and clearance (through nuclease action, liver, kidney, spleen, and immune function) mechanisms are complex and can influence biomarker detectability and persistence [65]. As such, tissue biomarkers are not by definition applicable for detection in LBs. Finally, APC is known to be methylated in gastrointestinal cancer types (no publications available on DPP7 methylation in cancer) [66]. As such, APC and (possibly also) DPP7 alone are likely not suitable for YST identification from a pan-cancer screening perspective, especially in older populations (where gastrointestinal cancers are more common) [67]. Importantly, combining both markers with miRNA-371a-3p detection could solve this issue.

We confirmed the previously identified collective GCT markers, *DPPA3* and *RASSF1A*, in our meta-analysis cohort [24, 25]. In contrast to *APC* and *DPP7, DPPA3* and *RASSF1A* are not YST-specific. Following DMR analysis for YST samples *versus* BNT and PB, *DPP7* and *APC* had a higher Δß (0.72 and 0.68) compared to *DPPA3* and *RASSF1A* (both 0.55) (data not shown). As such, *DPP7* and *APC* might have better sensitivity for YST detection (in tumor DNA) versus *DPPA3* and *RASSF1A*.

Our proof of concept encourages further identification and validation of other GCT subtype-specific biomarkers. Other GCT histologies include EC, TE, gGCTs (DG/GER/SE), CHC, ST, and DC. We identified multiple hypermethylated DMRs in EC and TE as potential biomarkers (Table 3). There, especially the TE group would be interesting to further explore as there is currently no true specific biomarker for it [68]. TEs are heterogeneous with well or incompletely differentiated somatic tissue, which makes finding a universal biomarker inherently difficult [1]. gGCTs are globally hypomethylated, which separates them from all other histologies. Importantly, our MSRE-qPCR approach is developed for detection of methylated DNA in a background of (digested) unmethylated DNA. As such, in the context of hypomethylation marker detection, unmethylated targets would be digested while background signal (methylated copies) will remain. Hence, this approach is more challenging, and possibly not feasible, especially in situations with low tumor signal with high background (e.g. LB). There are few reports that utilized methylation-dependent restriction enzymes (MDREs), such as MspJI, that only digest methylated DNA, and as such could be applied for detecting unmethylated DNA targets [69]. However, MDRE recognition sites are very specific and less abundant *versus* those of MSREs making amplicon selection and primer design more difficult (NEB website). Alternatively, a collectively methylated ngGCT marker (hence unmethylated in gGCTs and controls (BNT & PB)) could distinguish gGCTs from ngGCTs, as could global methylation profiling. Finally, while we did identify DMRs for CHC, ST, and DC, it should be noted that findings are less reliable *versus* DMRs for other histologies due to low sample size (3 or 4). Ultimately, development of a GCT subtype-specific biomarker panel would be relevant. The SYBR green-based MSRE-qPCR approach provides a fast and cheap way for screening potential markers for such a panel. For final panel design, however, a probe-based multiplex approach is preferred as SYBR green stains all double stranded DNA and is not target-specific. Such a multiplex approach would allow identification of multiple targets within a single sample, which is especially relevant in the context of LB where cfDNA availability is more limited *versus* tumor DNA. Also, an absolute quantitative setup, such as digital PCR (dPCR) would be preferred as this mitigates the need for inclusion of (target-specific) standard curves, further reducing labor intensity [70].

GCT (subtype-specific) biomarkers also enable developmental biology-orientated studies to elucidate GCT onset, pathogenesis, and progression. As example, *APC* is a negative regulator of the WNT signaling pathway, which is essential for internal homeostasis and frequently aberrantly activated in gastrointestinal cancer, notably through *APC* inactivation by mutation [66, 71]. YSTs are known to have active WNT signaling and high expression of WNT-related genes [72]. *APC* promotor hypermethylation in YSTs hypothetically supports this notion, given that promotor hypermethylation generally results in gene silencing, which would lead to reduced *APC* expression resulting in increased WNT signaling. However, a study by Okpanyi *et al* did not observe any changes in *APC* mRNA in YSTs *versus* other GCTs, while they did observe specific *APC* methylation in the YST group (11/13 cases) [73]. Concurrently, analysis of public GCT gene expression data using the R2 platform (identifier: ps_avgpres_gse3218gse10783ageo141_u133a) did not show differential gene expression in YSTs *versus* non-YSTs (Figure S27A). Therefore, other mechanisms beyond *APC* are likely involved in WNT activation. There are only very few other independent reports (with small sample sizes) investigating relations between YSTs and (epi)genetic aberrations of *APC* (mostly hypermethylation but also loss of heterozygosity and mutation), warranting the need for further studies [33, 74-76]. Our identification of significant WNT pathway enrichment based on aberrantly methylated genes exclusive to YST further supports the involvement of WNT signaling in YSTs. Interestingly, Xu *et al* showed that GCT CLs treatment with WNT inhibitors reduced CL growth especially in the YST CLs, suggesting opportunities for targeted treatment as WNT inhibitors are in clinical development for cancer treatment [33, 77].

*DPP7* is a member of the dipeptidyl peptidase family of proteins being important regulators of cellular metabolism and signaling cascades [78]. Specifically, the main function of *DPP7* is its role in maintaining survival of quiescent lymphocytes, while overexpression has been described in various tumors notably in colorectal cancer [78, 79]. Interestingly, using the same public GCT expression dataset as described above, we found that YSTs show significantly lower *DPP7* expression *versus* non-YSTs (Bonferroni-corrected Welch p < 0.01) (Figure 27B). Relations between *DPP7* and YST have not been described in the literature at all. As such, even though a direct relationship between DNA methylation and gene expression is not by definition relevant from a biomarker perspective, further investigating the link between *DPP7* promoter hypermethylation and reduced *DPP7* expression, and its role in YST onset and progression provides an interesting future direction of study.

We performed KEGG pathway enrichment on genes annotated to histology-specific DMRs to explore underlying biology. While gene sets were identified per subtype, substantial overlap was present across subtypes, resulting in largely shared pathways. YST was the only GCT subtype for which we identified enriched pathways based on differentially methylated genes exclusive to this subtype. This indicates the limited information provided by methylation data alone and highlights the need for gene expression data to better understand subtype-specific gene regulation in GCTs.

Despite its comprehensiveness, our study has some limitations. Firstly, we did not include any healthy ovary and brain tissue in our meta-analysis as additional controls next to healthy testis and peripheral blood. GCTs can also occur in the ovary and brain, and potential methylation in these healthy tissues is currently not considered in our meta-analysis. Using the EWAS platform, we found that probes covering our *APC* and *DPP7* amplicons are not methylated in healthy brain (ovary data was not available) (data not shown) [64]. Also, our meta-analysis is based on tumor tissue DNA where findings are not always transferrable to application in LB. For specific LB biomarker discovery, 450K/EPIC arrays are likely not optimal as they require 10-500 ng input DNA (which can exceed available cfDNA), are limited in CpG coverage, and might mask tumor signal by background non-tumor-derived cfDNA signal [80]. There are several alternative sulfite, enzyme, or binding-based methods available that can be tailored for cfDNA-based methylation profiling [80]. Nonetheless, GCT-specific cfDNA-based methylation data is scarce, likely due to the field having shifted towards microRNAs [81]. As such, our meta-analysis still provides a comprehensive platform for identification of potential LB-based biomarkers, proven by *DPP7*.

In conclusion, our meta-analysis indicates that GCT histologies have distinct methylation profiles that can be utilized for subtype-specific biomarker identification and validation using our bioinformatic and MSRE-qPCR pipeline. Subtype-specific GCT biomarkers could complement or outperform the classical serum markers and together with miRNA-371a-3p provide a sensitive and specific companion diagnostic for informing GCT management during different stages of care. To our knowledge, *DPP7* represents the first YST-specific methylation-based cfDNA biomarker. Future validation in larger independent cohorts and longitudinal monitoring throughout different clinical stages is needed to confirm clinical utility. Finally, our bioinformatic pipeline is easily transferrable. This encourages similar applications in pan(pediatric)-cancer studies beyond GCTs.

## Supporting information

Supplemental results, figures, and table legends

## Data availability

Original DNA methylation array data (datasets 2, 3, 7) has been submitted to the NCBI Gene Expression Omnibus (GEO) under accession numbers GSE303824, GSE303824, and GSE303824, respectively.

## Code availability

The final developed bioinformatic pipeline for GCT methylation meta-analysis is provided as script (supplemental Rscript).

## Contributions

Conceptualization: F.J. and L.L.; writing and first draft preparation: F.J.; design of bioinformatic meta-analysis pipeline: F.J., L.K., T.E.; bioinformatic meta-analysis: F.J.; wet lab work: F.J., A.G.; revisions and editing: F.J., A.G., L.K, H.K., K.I., T.E., L.L.; supervision: L.L.; corresponding author: L.L. All authors read and approved the final manuscript.

## Acknowledgements

The authors would like to thank Marnix van Soest for culturing and provision of GCT-44 and NOY-1 CLs. This study was funded by the Dutch foundation Stichting Kinderen Kankervrij (KiKa). The funder was not involved in the study nor writing of this manuscript.

