## Supplemental results, figures, and table legends for "Identification and validation of liquid biopsy-based methylation biomarkers: a germ cell tumor subtype-specific study"

Several primer pairs were designed for different amplicon targets within the DMRs for *APC* and *DPP7*. Specificity was investigated using gel electrophoresis following endpoint PCR and melting curves following qPCR (Figure S17). Per target, the best primer pair was chosen (Table S2 for sequences). Assay development experiments were conducted using the GCT CLs TCam-2 and JEG, which were respectively hypo and hypermethylated for both targets based on our methylation meta-analysis dataset (Figure S18). Sufficient MSRE input is required for full digestion of unmethylated DNA. However, excess MSRE input could result in undesired nonspecific digestion of methylated DNA. Hence, careful optimization of these parameters is required. Both targets had (multiple) restriction sites for different enzymes (*APC*: 3 for AciI, 2 for HhaI, 1 for Fnu4HI. *DPP7*: 1 for AciI, 1 for HhaI, 2 for HpaII, 1 for Fnu4HI, and 1 for AfeI). Hence, we started with testing different enzyme combinations. A combination of AciI and HhaI resulted in better *APC* digestion *versus* individual enzymes (Figure S19). For *DPP7*, a combination of HpaII and HhaI also resulted in better digestion, which was not improved by adding a third enzyme (Figure S19). Also, enzyme inactivation after the digestion step is essential to prevent additional digestion during the PCR steps. 10 minutes of incubation at 95°C was sufficient to inactivate activity for all enzymes except Fnu4HI, hence we refrained from further using this enzyme at all (Figure S20). Using these enzyme combinations, we investigated a wide range of enzyme and DNA input for ratio optimization. For both targets, 0.1 (total) enzyme units/ng DNA input was deemed optimal. A higher ratio did not improve TCam-2 digestion but did result in JEG-3 digestion (only for *APC*) indicating aspecific digestion of methylated DNA (Figure 5A). For *DPP7*, increasing the ratio could possibly result in slightly improved TCam-2 digestion, however, this difference was not big enough to justify the exponential increase in enzyme usage (Figure 5B). The defined 0.1 ratio consisted of 1 unit per individual enzyme (hence 2 in total) and 20ng input DNA. 15 and 60 minutes of enzyme incubation using these conditions resulted in similar digestion efficacy, hence 15 minutes was chosen for saving time (data not shown). To improve practicality and save time, we performed all digestion steps in the qPCR mix (hence no cleanup step in between). NEB recommended using their rCutSmart Buffer for optimal digestion, however, we found that it inhibited the qPCR reaction resulting in no target amplification (data not shown). Hence, we refrained from further using this buffer. Following these optimization steps, we tested different GCT CLs (TCam-2, NCCIT, 2102-EP, NT-2, GCT-44, NOY-1, JEG-3, JAR), DNA from a healthy male and female donor, and commercially available methylated and unmethylated DNA. For the *APC* assay, GCT-44 and NOY-1 were selected as final methylated and unmethylated controls. For the *DPP7* assay, JAR and NOY-1 were selected as controls. These CLs had the highest and lowest ΔCt thus representing the best unmethylated and methylated conditions, respectively, available to us (Figure 5C). Also, these samples performed similar (or better) than commercially available methylated and unmethylated DNA, and were therefore chosen to save costs. For both assays, standard curves correlating ΔCt and methylation % (where the methylated and unmethylated controls were expressed as 100% and 0% methylated, respectively) were generated (Figure 5D). We measured a similar methylation % in conditioned medium of GCT-44 as in GCT-44 cells (not tested for other GCT CLs), indicating that our assays could be suitable for methylation monitoring during GCT CL culturing experiments (Figure S21).

### Supplementary Figures

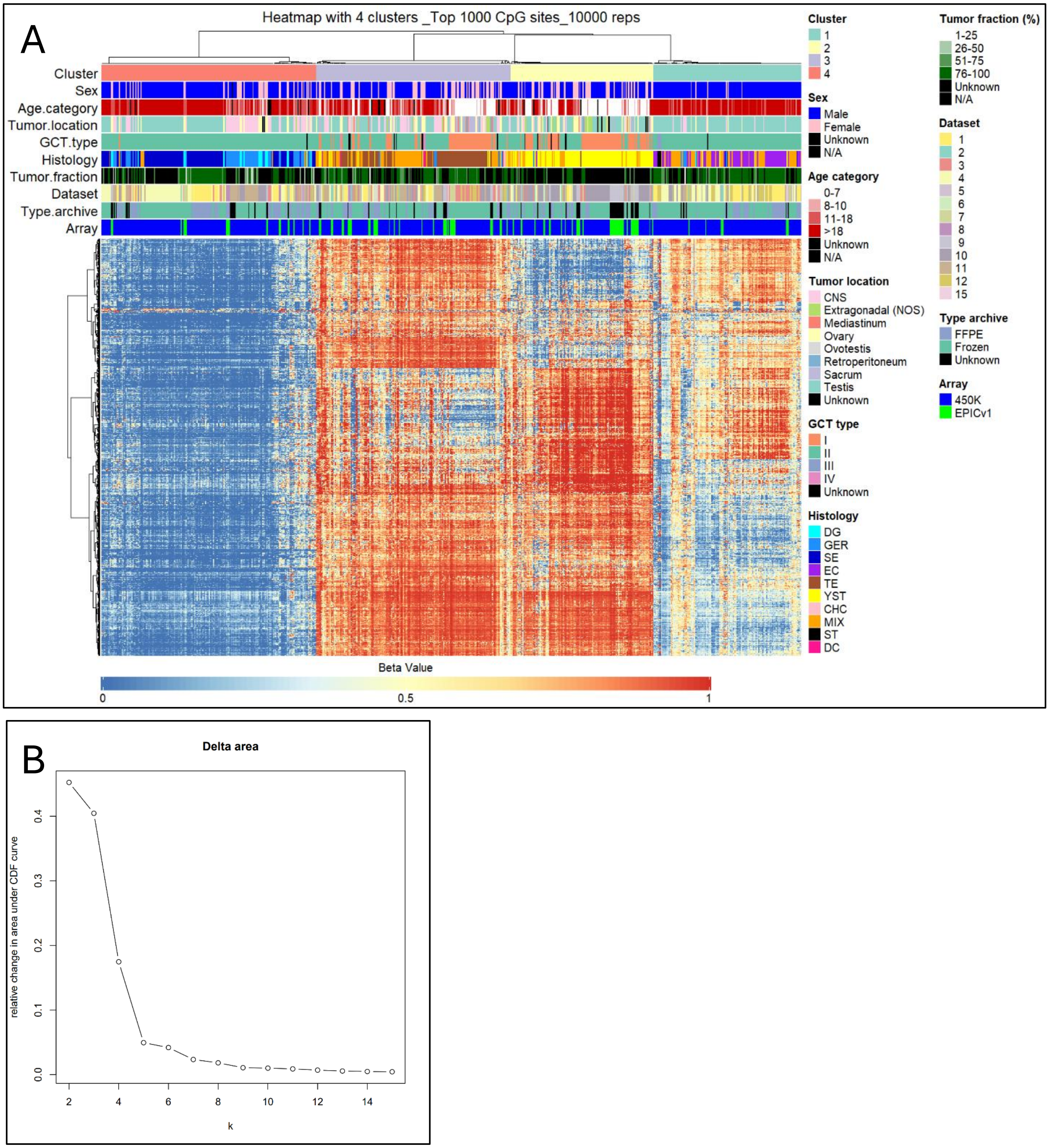


***Figure S1: inclusion of mixed GCT samples maintains clustering of GCTs with the same pure histology regardless of other parameters****. Methylation-based UHC of GCT samples with pure histology + mixed samples was performed on beta-values, using the top 1000 CpG sites with highest median absolute variation and 10,000 repetitions (A). Heatmap columns represent unique samples, rows represent unique CpG sites. Core data represents beta-values ranging from 0 (fully demethylated) to 1 (fully methylated) based on a color gradient scale. Annotations with several clinical and histological parameters are provided on top with corresponding legends provided on the right. The amount of clusters was chosen based on visual inspection of the corresponding Delta area plot (B). There, the relative change in area under the Consensus Cumulative Distribution Function (CDF) curve is presented comparing k (number of clusters) and k -1. This allows determination of the relative increase in consensus and determine k at which there is no appreciable increase. In this specific graph, k = 4 was chosen. Abbreviations: CHC: choriocarcinoma, DC: dermoid cyst, DG: dysgerminoma, EC: embryonal carcinoma, GER: germinoma, MIX: GCT with mixed histologies, SE: seminoma, ST: spermatocytic tumor, TE: teratoma, YST: yolk sac tumor.*

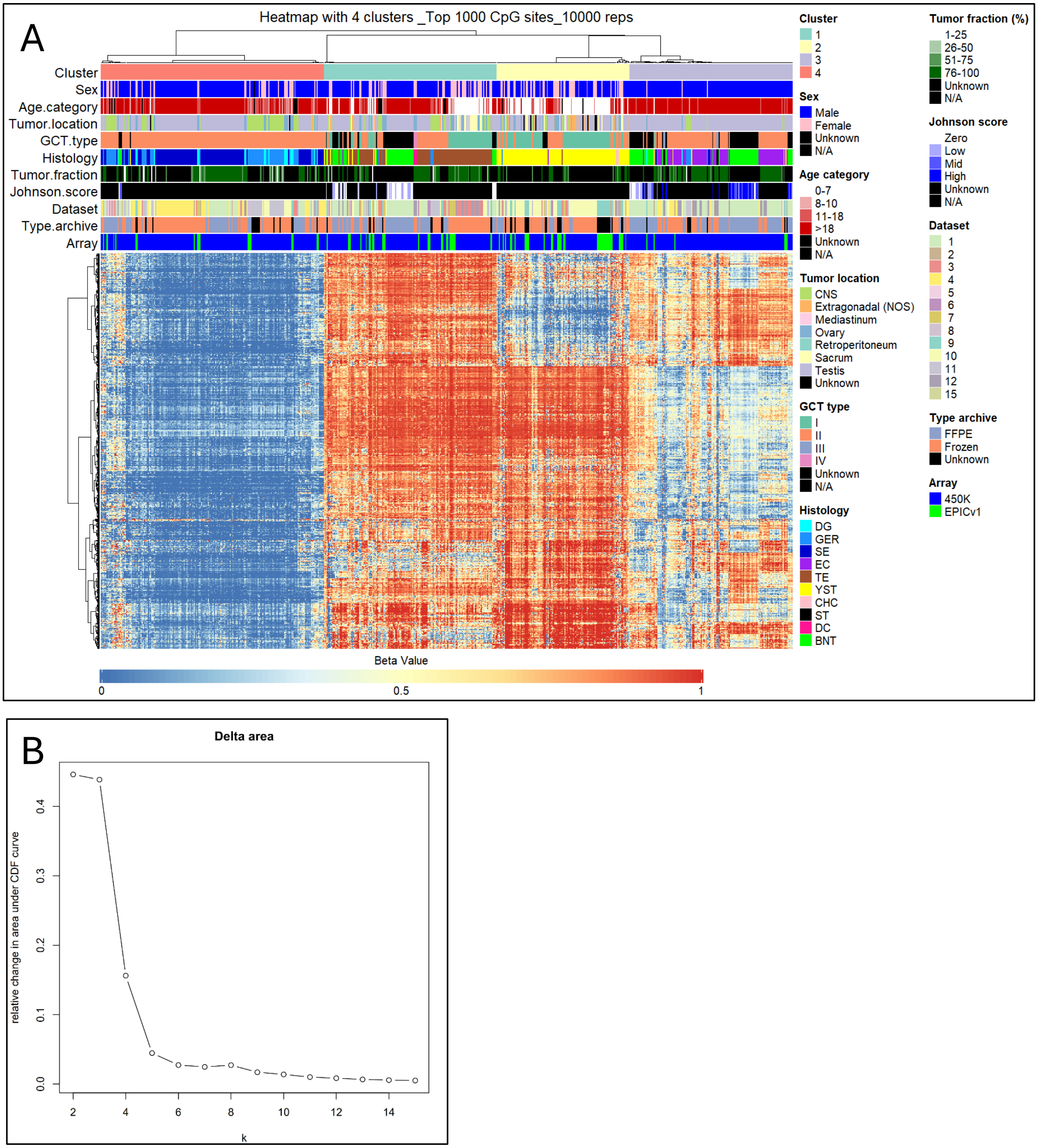


***Figure S2: BNT samples cluster partly with TE and partly with EC samples***. *Methylation-based UHC of GCT samples with pure histology + BNT samples was performed on beta-values, using the top 1000 CpG sites with highest median absolute variation and 10,000 repetitions (A). Heatmap columns represent unique samples, rows represent unique CpG sites. Core data represents beta-values ranging from 0 (fully demethylated) to 1 (fully methylated) based on a color gradient scale. Annotations with several clinical and histological parameters are provided on top with corresponding legends provided on the right. The amount of clusters was chosen based on visual inspection of the corresponding Delta area plot (B). There, the relative change in area under the Consensus Cumulative Distribution Function (CDF) curve is presented comparing k (number of clusters) and k -1. This allows determination of the relative increase in consensus and determine k at which there is no appreciable increase. In this specific graph, k = 4 was chosen. Abbreviations: BNT: benign neighboring testis, CHC: choriocarcinoma, DC: dermoid cyst, DG: dysgerminoma, EC: embryonal carcinoma, GER: germinoma, SE: seminoma, ST: spermatocytic tumor, TE: teratoma, YST: yolk sac tumor*

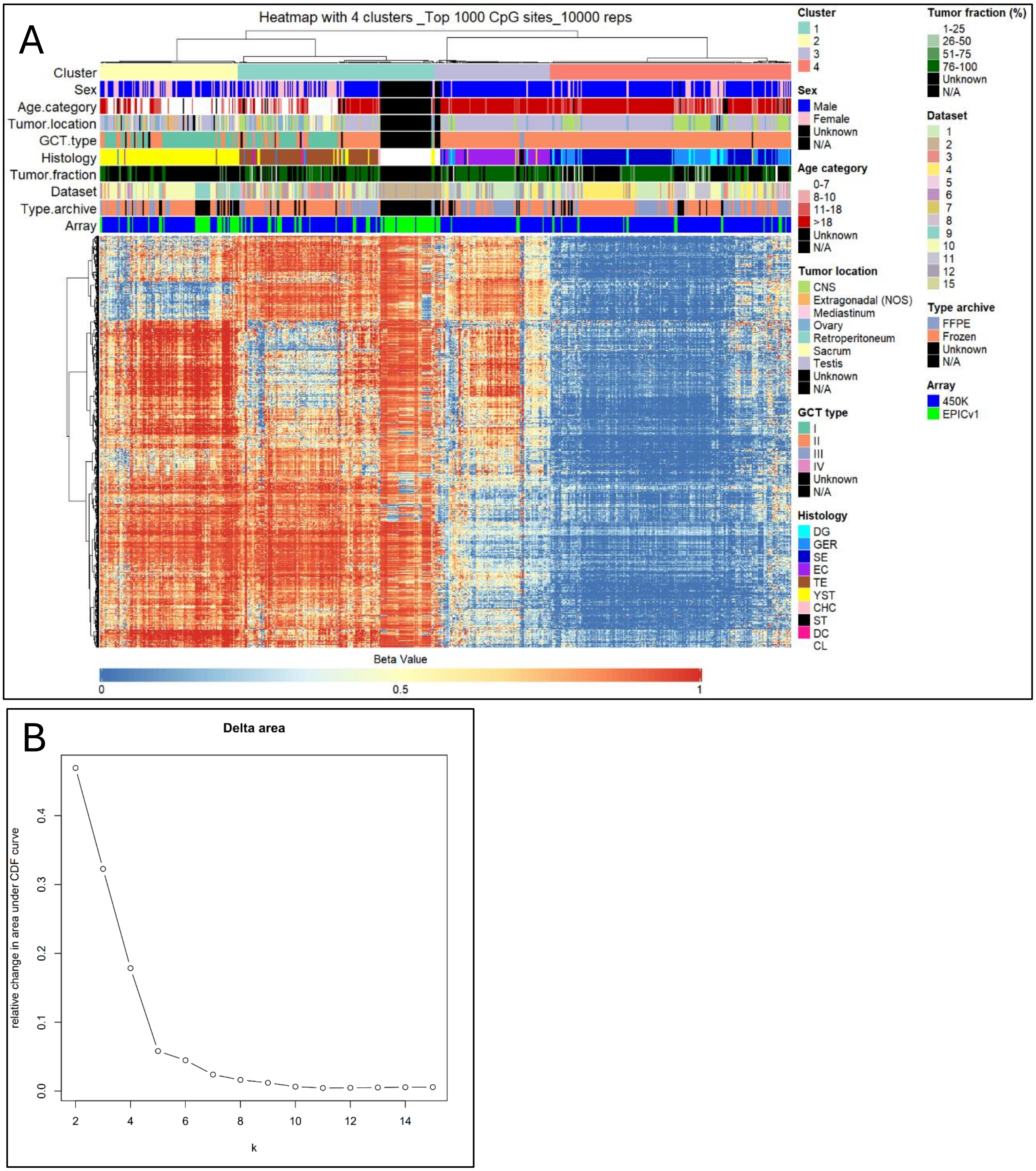


***Figure S3: GCT CLs cluster with TE samples**. Methylation-based UHC of GCT samples with pure histology + GCT CLs was performed on beta-values, using the top 1000 CpG sites with highest median absolute variation and 10,000 repetitions (A). Heatmap columns represent unique samples, rows represent unique CpG sites. Core data represents beta-values ranging from 0 (fully demethylated) to 1 (fully methylated) based on a color gradient scale. Annotations with several clinical and histological parameters are provided on top with corresponding legends provided on the right. The amount of clusters was chosen based on visual inspection of the corresponding Delta area plot (B). There the relative change in area under the Consensus Cumulative Distribution Function (CDF) curve is presented comparing k (number of clusters) and k -1. This allows determination of the relative increase in consensus and determine k at which there is no appreciable increase. In this specific graph, k = 4 was chosen. Abbreviations: CL: cell line, CHC: choriocarcinoma, DC: dermoid cyst, DG: dysgerminoma, EC: embryonal carcinoma, GER: germinoma, SE: seminoma, ST: spermatocytic tumor, TE: teratoma, YST: yolk sac tumor*

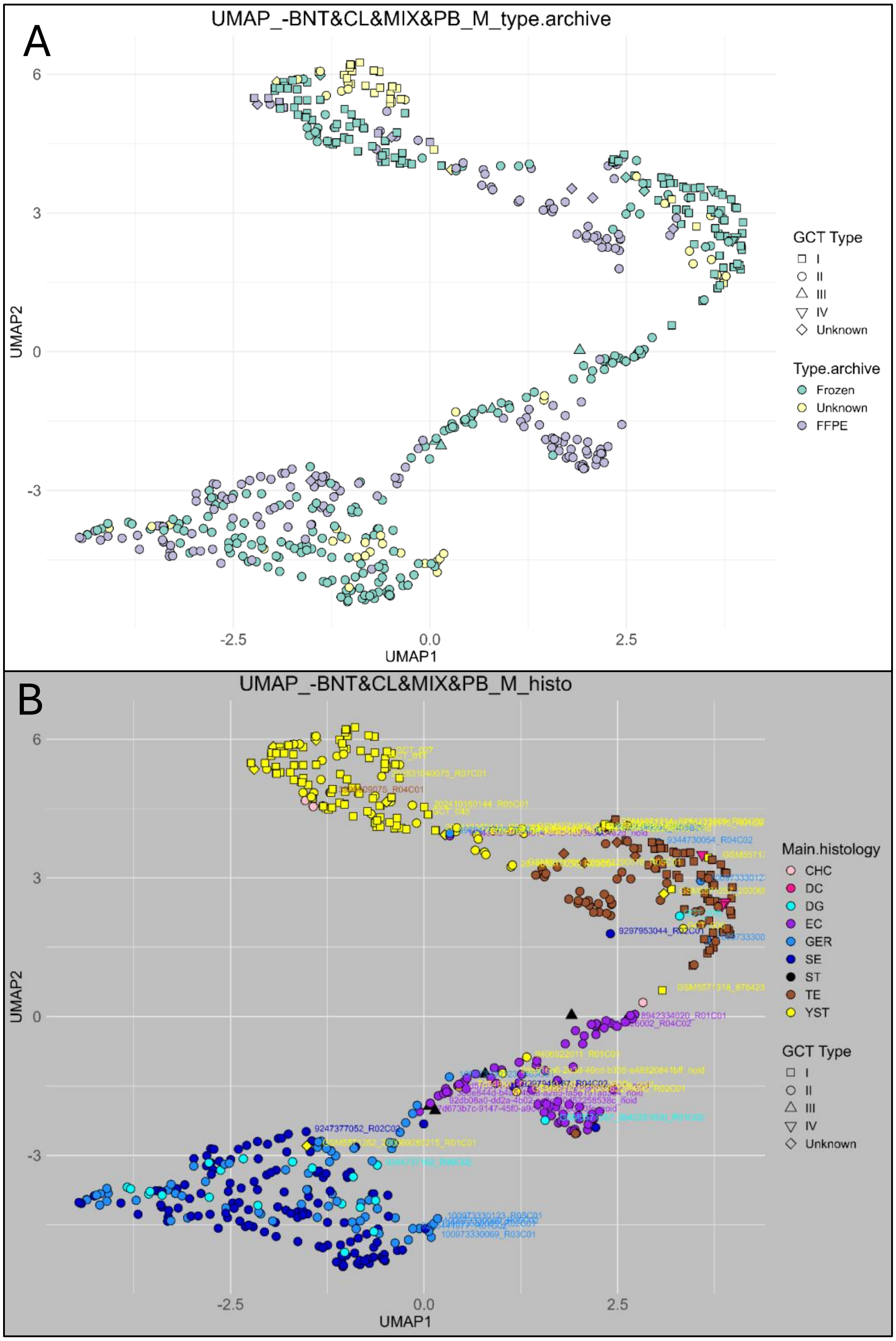


***Figure S4: UMAP with only GCT samples of pure histology***. *Data points are identical to those presented in main Figure 3. For A: data point color refers to the tissue archive type (FFPE or frozen). For B: data point color refers to histology. Highlighted samples (with sample IDs next to their corresponding data points) indicate samples that clustered outside of their respective main cluster based on Figure 2. For both A and B, data point shape indicates GCT type (relevant only for YST and TE samples). Abbreviations: CHC: choriocarcinoma, DC: dermoid cyst, DG: dysgerminoma, EC: embryonal carcinoma, GER: germinoma, SE: seminoma, ST: spermatocytic tumor, TE: teratoma, YST: yolk sac tumor.*

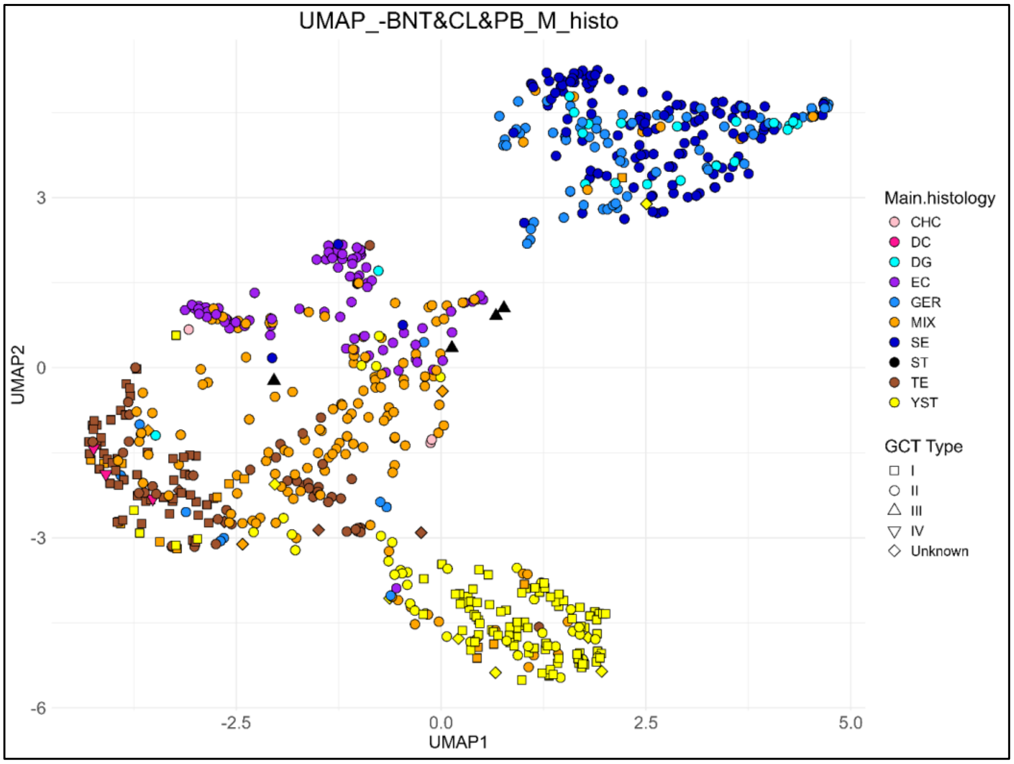


***Figure S5: Mixed GCT samples group together within and between individual GCT samples of pure histology**. UMAP of GCT samples with pure histology + mixed samples was performed on M-values. Samples are color-coded by their histology. Data point shapes refer to GCT type (relevant only for YST and TE samples). Abbreviations: CHC: choriocarcinoma, DC: dermoid cyst, DG: dysgerminoma, EC: embryonal carcinoma, GER: germinoma, MIX: GCT with mixed histologies, SE: seminoma, ST: spermatocytic tumor, TE: teratoma, YST: yolk sac tumor.*

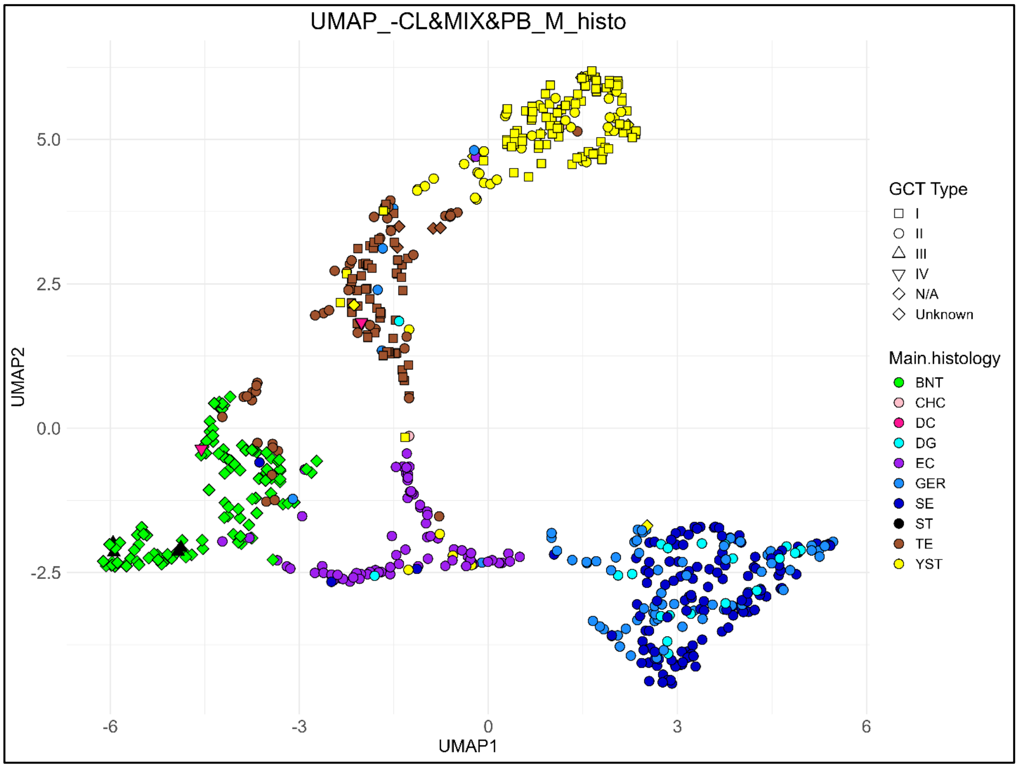


***Figure S6: BNTs form a distinct group together with a subset of TE samples**. UMAP of GCT samples with pure histology + BNT was performed on M-values. Samples are color-coded by their histology. Data point shapes refer to GCT type (relevant only for YST and TE samples). Abbreviations: BNT: benign neighboring testis, CHC: choriocarcinoma, DC: dermoid cyst, DG: dysgerminoma, EC: embryonal carcinoma, GER: germinoma, SE: seminoma, ST: spermatocytic tumor, TE: teratoma, YST: yolk sac tumor.*

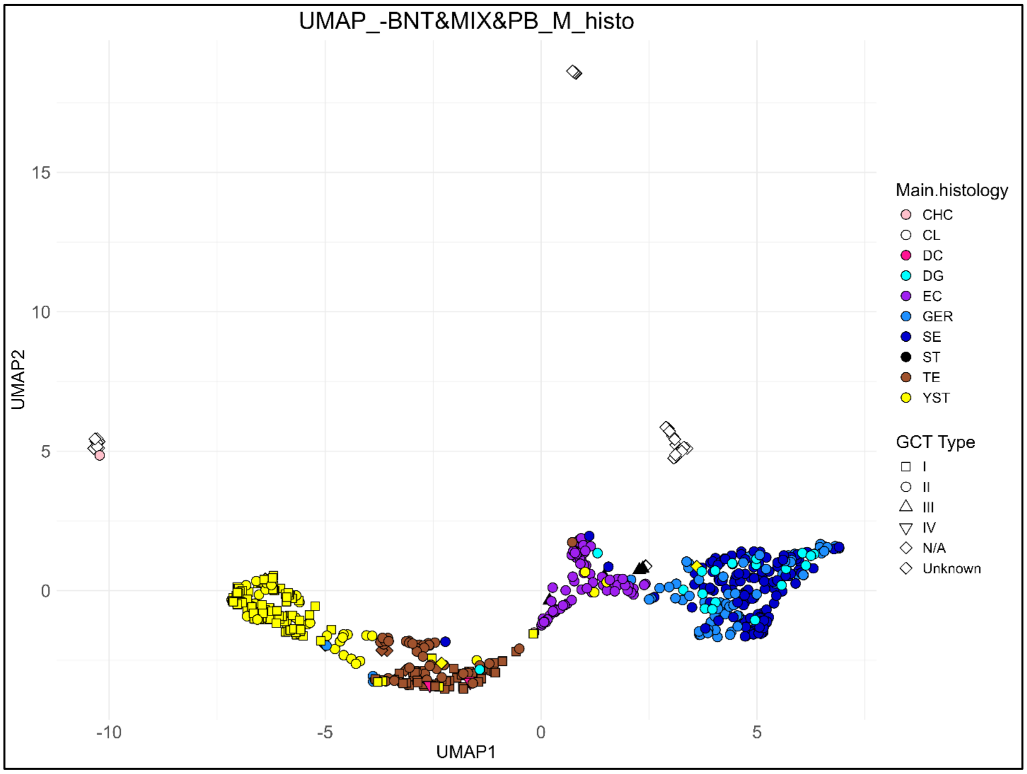


***Figure S7: GCT CLs form distinct groups completely separated from GCT samples**. UMAP of GCT samples with pure histology + GCT CLs was performed on M-values. Samples are color-coded by their histology. Data point shapes refer to GCT type (relevant only for YST and TE samples). Abbreviations: CL: cell line, CHC: choriocarcinoma, DC: dermoid cyst, DG: dysgerminoma, EC: embryonal carcinoma, GER: germinoma, SE: seminoma, ST: spermatocytic tumor, TE: teratoma, YST: yolk sac tumor.*

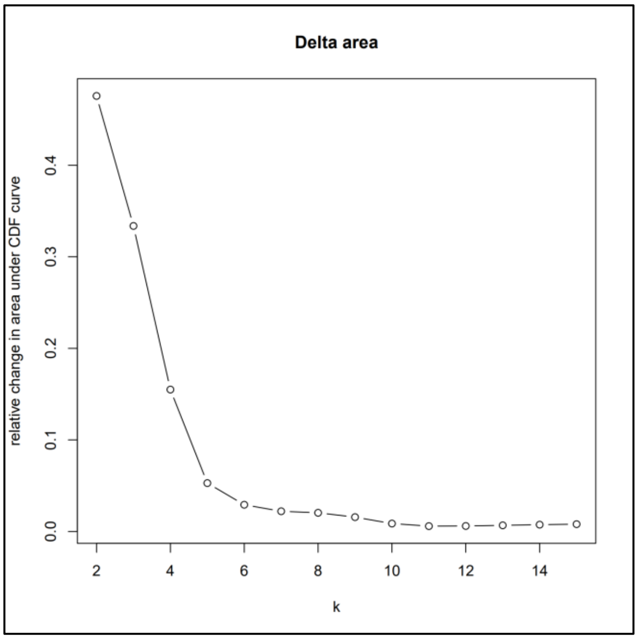


***Figure S8: delta area plot of UHC containing only pure GCT samples.*** The relative change in area under the Consensus Cumulative Distribution Function (CDF) curve is presented comparing k (number of clusters) and k -1. This allows determination of the relative increase in consensus and determine k at which there is no appreciable increase. This delta area plot is linked to the data and heatmap visualized in Figure 2. In this specific graph, k = 4 was chosen.

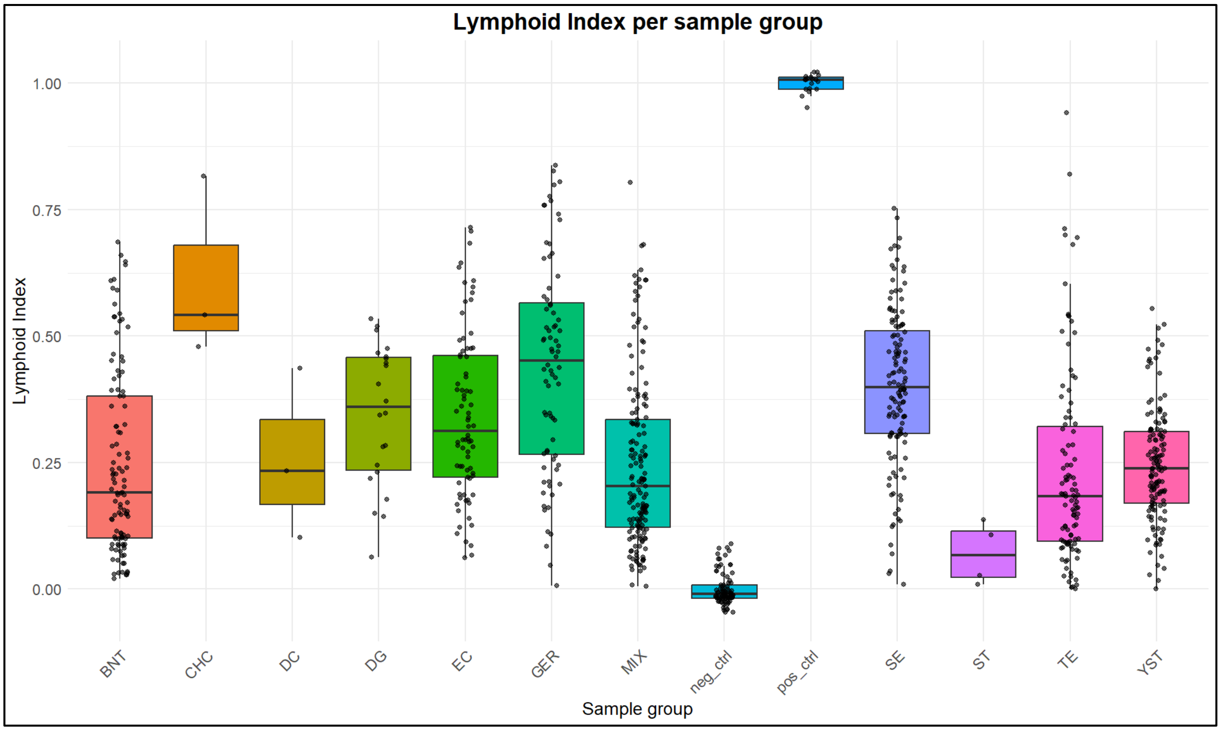


***Figure S9: GCT tissues can display varying amounts of lymphocyte infiltration***. *Lymphoid index (LI) is displayed per sample group. A sample's LI was defined by taking the average beta-value of a set of 36 specific probes. Details for LI calculation are described in the methods section. Abbreviations: BNT: benign neighboring testis, CHC: choriocarcinoma, DC: dermoid cyst, DG: dysgerminoma, EC: embryonal carcinoma, GER: germinoma, MIX: GCT with mixed histologies, SE: seminoma, ST: spermatocytic tumor, TE: teratoma, YST: yolk sac tumor.*

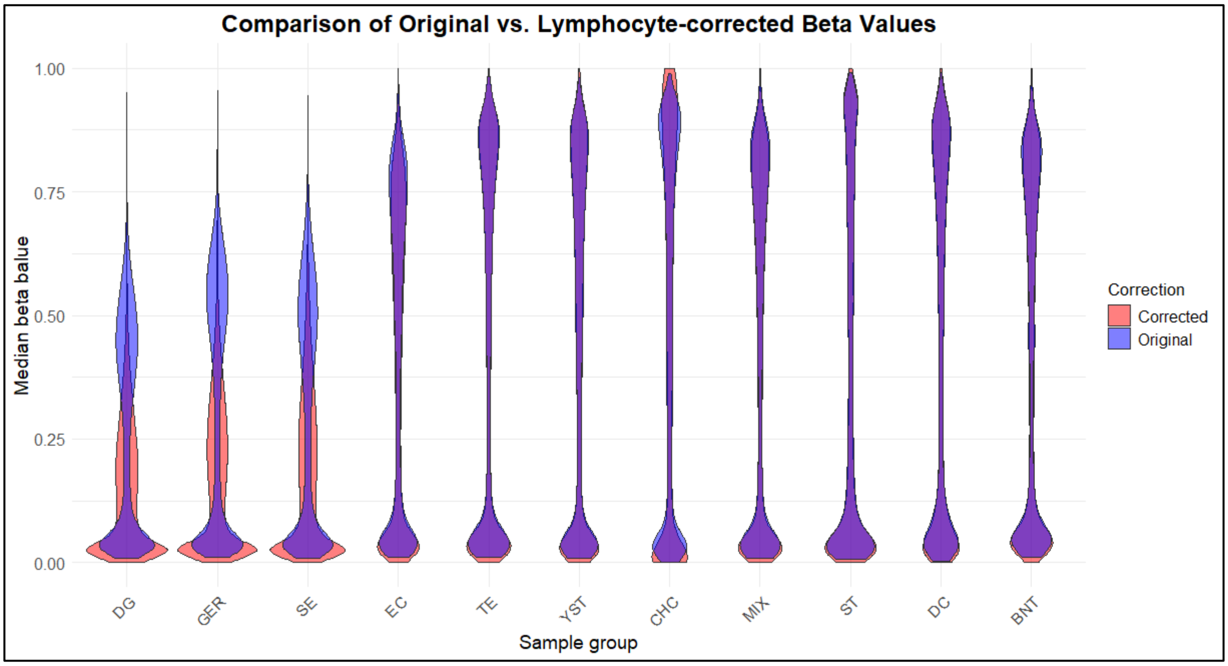


***Figure S10: correction for lymphocyte infiltration results in a reduction of the global methylation profile for gGCTs.*** *The median beta value distribution harmonized for all samples per group (see methods for details) are displayed. Color coding indicates global methylation profiles before (blue) and after (red) correction. Abbreviations: BNT: benign neighboring testis, CHC: choriocarcinoma, DC: dermoid cyst, DG: dysgerminoma, EC: embryonal carcinoma, GER: germinoma, MIX: GCT with mixed histologies, SE: seminoma, ST: spermatocytic tumor, TE: teratoma, YST: yolk sac tumor.*

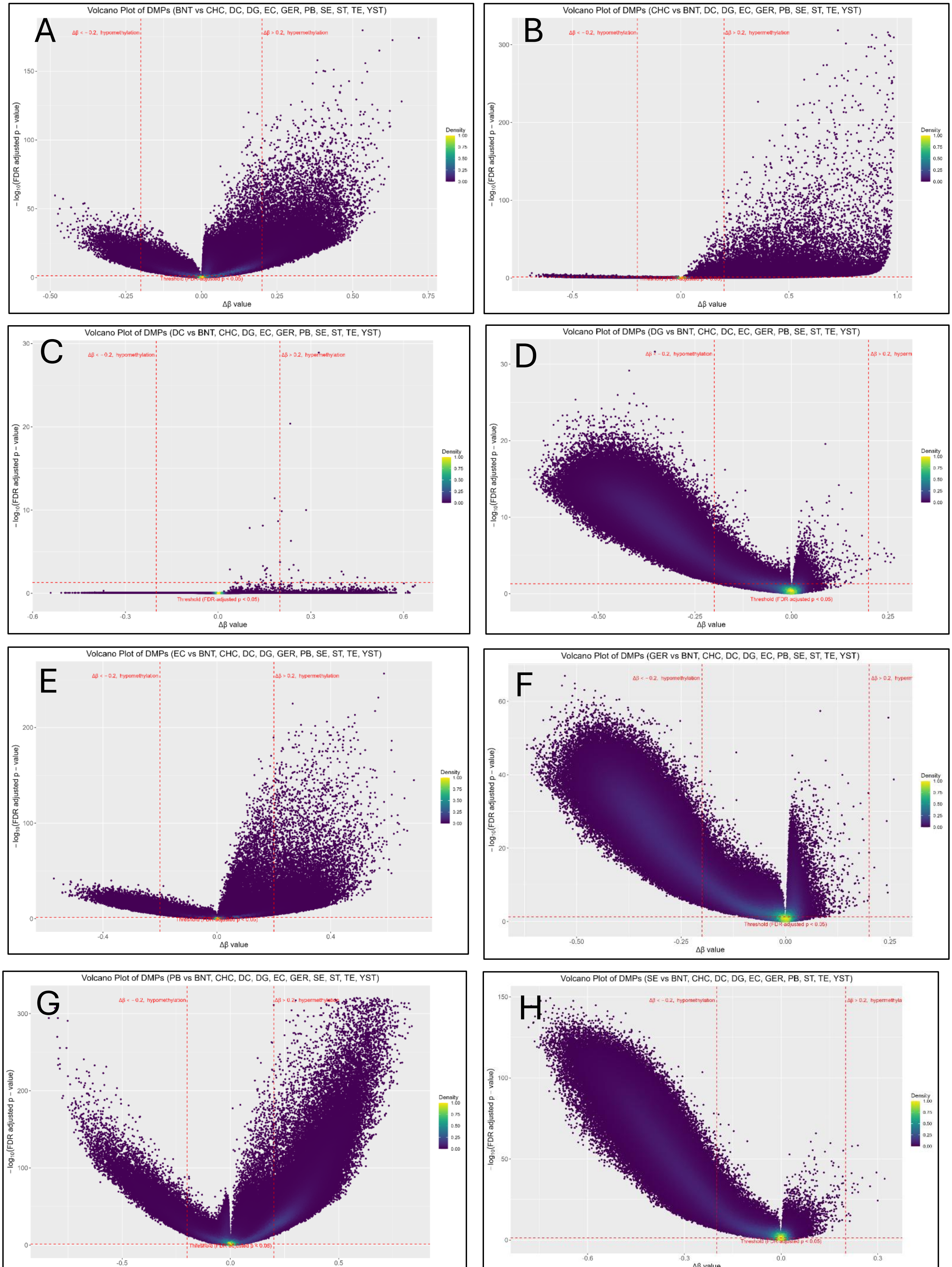

Volcano Plot of DMPs (BNT vs CHC, DC, DG, EC, GER, PB, SE, ST, TE, YST)
A
Δβ < −0.2, hypomethylation
Δβ > 0.2, hypermethylation
Threshold (FDR-adjusted p < 0.05)
− log10(FDR adjusted p − value)
Δβ value
Density
Volcano Plot of DMPs (CHC vs BNT, DC, DG, EC, GER, PB, SE, ST, TE, YST)
B
Δβ < −0.2, hypomethylation
Δβ > 0.2, hypermethylation
− log10(FDR adjusted p − value)
Δβ value
Density
Volcano Plot of DMPs (DC vs BNT, CHC, DG, EC, GER, PB, SE, ST, TE, YST)
C
Δβ < −0.2, hypomethylation
Δβ > 0.2, hypermethylation
Threshold (FDR-adjusted p < 0.05)
− log10(FDR adjusted p − value)
Δβ value
Density
Volcano Plot of DMPs (DG vs BNT, CHC, DC, EC, GER, PB, SE, ST, TE, YST)
D
Δβ < −0.2, hypomethylation
Δβ > 0.2, hyperm
Threshold (FDR-adjusted p < 0.05)
− log10(FDR adjusted p − value)
Δβ value
Density
Volcano Plot of DMPs (EC vs BNT, CHC, DC, DG, GER, PB, SE, ST, TE, YST)
E
Δβ < −0.2, hypomethylation
Δβ > 0.2, hypermethylation
− log10(FDR adjusted p − value)
Δβ value
Density
Volcano Plot of DMPs (GER vs BNT, CHC, DC, DG, EC, PB, SE, ST, TE, YST)
F
Δβ < −0.2, hypomethylation
Δβ > 0.2, hyperm
Threshold (FDR-adjusted p < 0.05)
− log10(FDR adjusted p − value)
Δβ value
Density
Volcano Plot of DMPs (PB vs BNT, CHC, DC, DG, EC, GER, SE, ST, TE, YST)
G
Δβ < −0.2, hypomethylation
− log10(FDR adjusted p − value)
Density
Volcano Plot of DMPs (SE vs BNT, CHC, DC, DG, EC, GER, PB, ST, TE, YST)
H
Δβ < −0.2, hypomethylation
Δβ > 0.2, hypermethyla
Threshold (FDR-adjusted p < 0.05)
− log10(FDR adjusted p − value)
Density

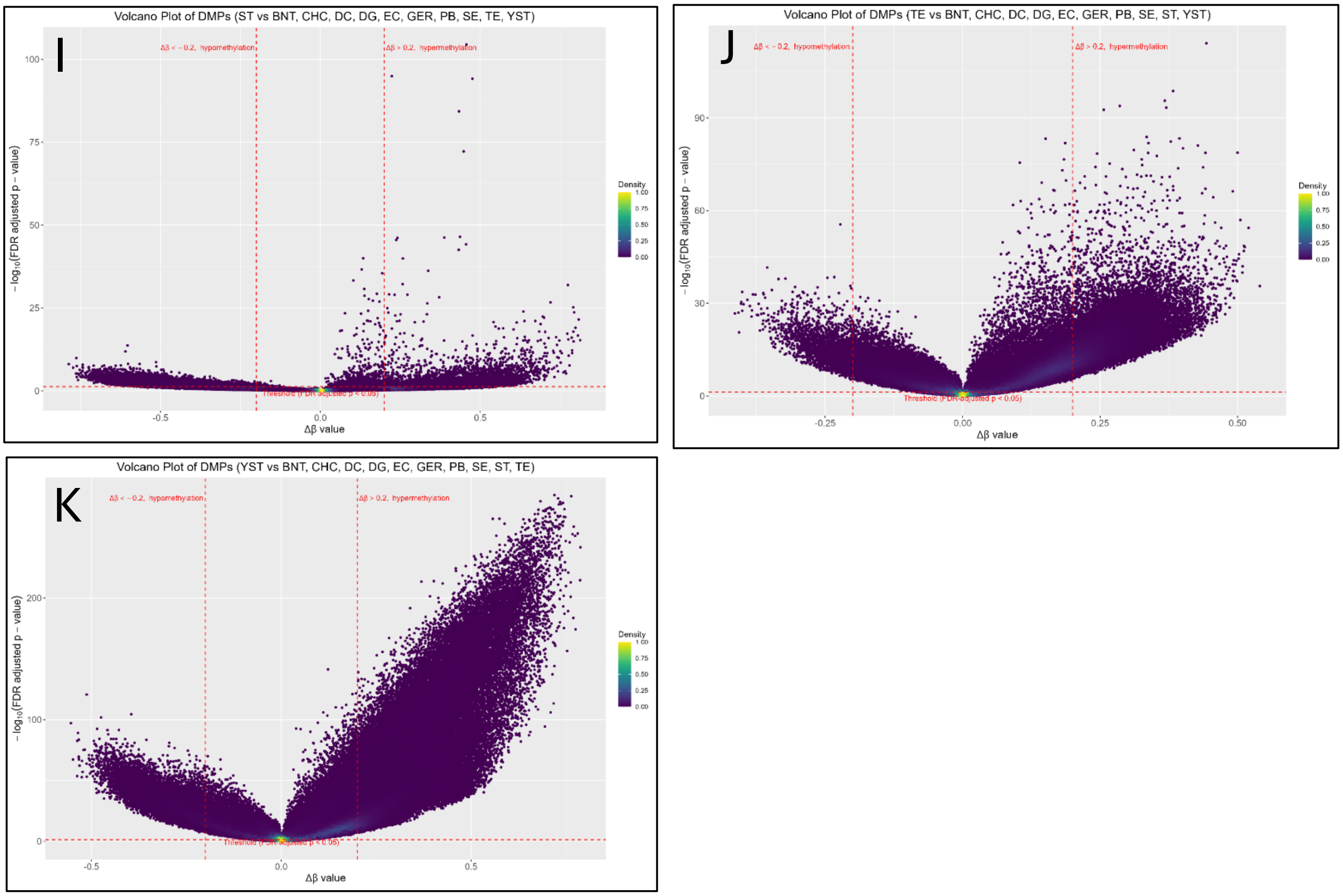


***Figure S11: Volcano plot with sample group-specific DMPs.*** *DMPs specific for each sample group: BTN (A), CHC (B), DC (C), DG (D), EC (E), GER (F), PB (G), SE (H), ST (I), TE (J), YST (K) were calculated (see methods for details). Per DMP/ probe/CpG site, -log10(FDR-adjusted p-values) and delta beta-values are presented. Color coding indicates density of overlapping probes. Thresholds for hypermethylated and hypomethylated probes, and significance are indicated by red lines. Abbreviations: DMP: differentially methylated position. BNT: benign neighboring testis, CHC: choriocarcinoma, DC: dermoid cyst, DG: dysgerminoma, EC: embryonal carcinoma, GER: germinoma, PB: peripheral blood, SE: seminoma, ST: spermatocytic tumor, TE: teratoma, YST: yolk sac tumor, FDR: false detection rate.*

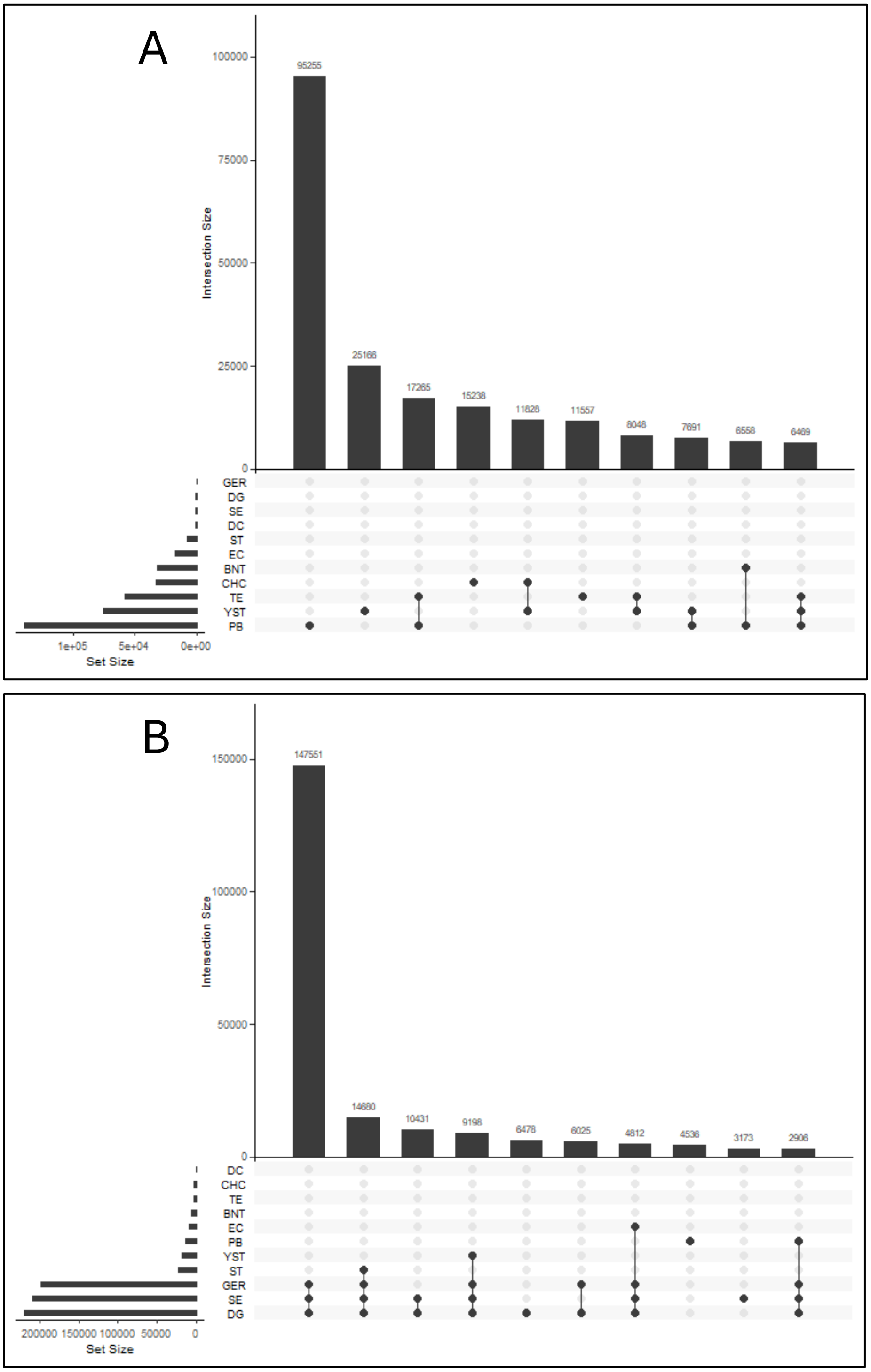


***Figure S12: collectively shared hypermethylated (A) and hypomethylated (B) probes among GCT histologies, BNT, and PB.*** *Sample group specific DMPs/probes/CpG sites (Figure S11) were collected and overlapping DMPs were identified using Upset plotting. Intersection Size=total hyper/hypomethylated probe number in a single group, or being shared among multiple groups (indicated by bullet points below that column). Set Size=total number of hyper/hypo methylated probes per group regardless of presence in other groups. Abbreviations: BNT: benign neighboring testis, CHC: choriocarcinoma, DC: dermoid cyst, DG: dysgerminoma, EC: embryonal carcinoma, GER: germinoma, PB: peripheral blood, SE: seminoma, ST: spermatocytic tumor, TE: teratoma, YST: yolk sac tumor.*

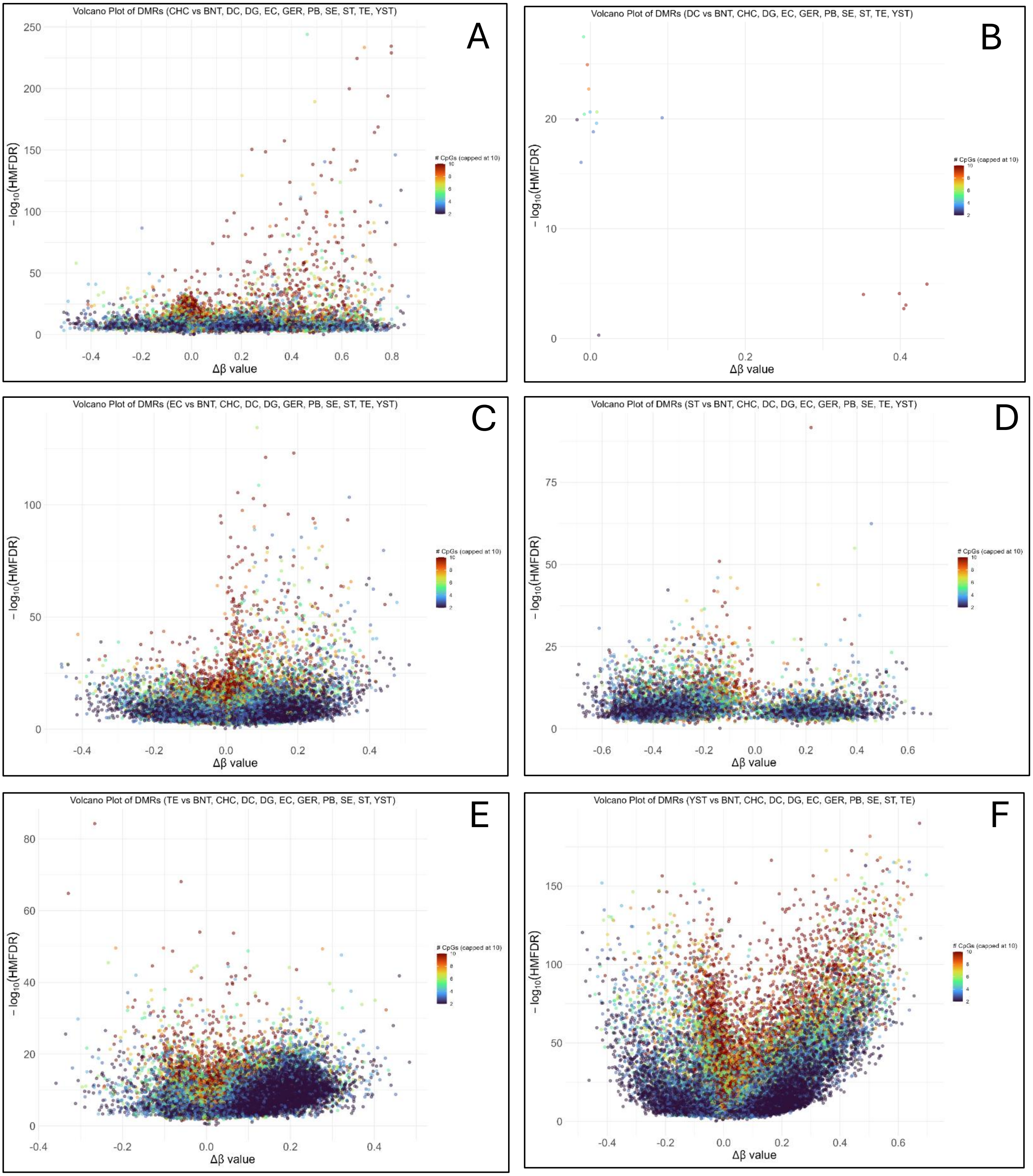
Volcano Plot of DMRs (CHC vs BNT, DC, DG, EC, GER, PB, SE, ST, TE, YST)
A
− log$_{10}$(HMFDR)
Δβ value
### CpGs (capped at 10)
Volcano Plot of DMRs (DC vs BNT, CHC, DG, EC, GER, PB, SE, ST, TE, YST)
B
− log$_{10}$(HMFDR)
Δβ value
### CpGs (capped at 10)
Volcano Plot of DMRs (EC vs BNT, CHC, DC, DG, GER, PB, SE, ST, TE, YST)
C
− log$_{10}$(HMFDR)
Δβ value
### CpGs (capped at 10)
Volcano Plot of DMRs (ST vs BNT, CHC, DC, DG, EC, GER, PB, SE, TE, YST)
D
− log$_{10}$(HMFDR)
Δβ value
### CpGs (capped at 10)
Volcano Plot of DMRs (TE vs BNT, CHC, DC, DG, EC, GER, PB, SE, ST, YST)
E
− log$_{10}$(HMFDR)
Δβ value
### CpGs (capped at 10)
Volcano Plot of DMRs (YST vs BNT, CHC, DC, DG, EC, GER, PB, SE, ST, TE)
F
− log$_{10}$(HMFDR)
Δβ value
### CpGs (capped at 10)

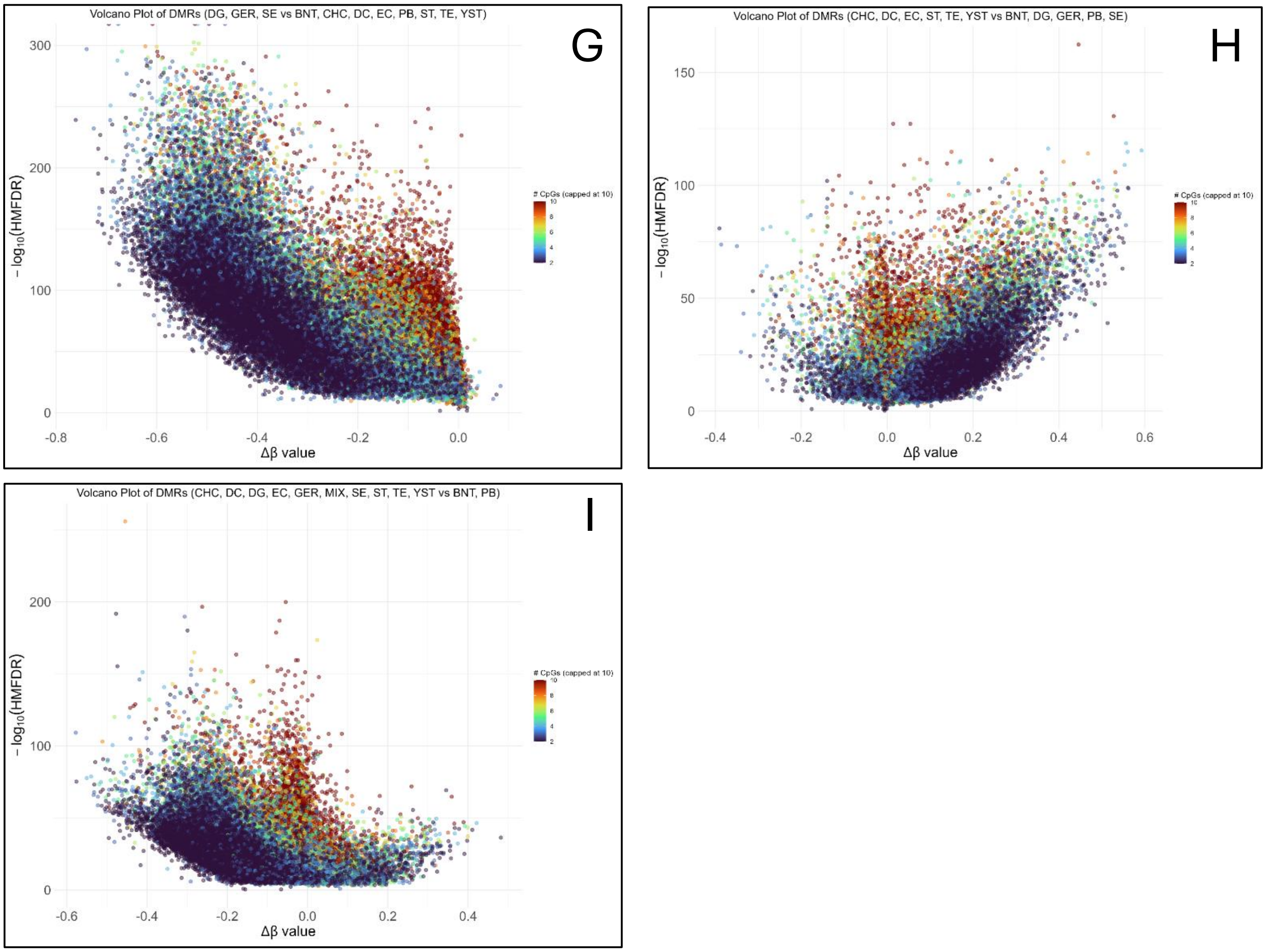


***Figure S13: Volcano plot with sample group-specific DMRs.*** *DMRs specific to individual sample groups: CHC (A), DC (B), EC (C), ST (D), TE (E), YST (F), as well as collective gGCTs (G), ngGCTs (H), and all GCTs (I) were calculated (see methods for details). Per DMR/, -log10(HMFDR) and delta beta-values are presented. Color coding indicates the number of CpG sites/probes per DMR. Abbreviations: DMP: differentially methylated position. BNT: benign neighboring testis, CHC: choriocarcinoma, DC: dermoid cyst, DG: dysgerminoma, EC: embryonal carcinoma, GER: germinoma, PB: peripheral blood, SE: seminoma, ST: spermatocytic tumor, TE: teratoma, YST: yolk sac tumor, HMFDR: harmonic mean of the FDR-corrected individual CpG p-values that make up a DMR.*

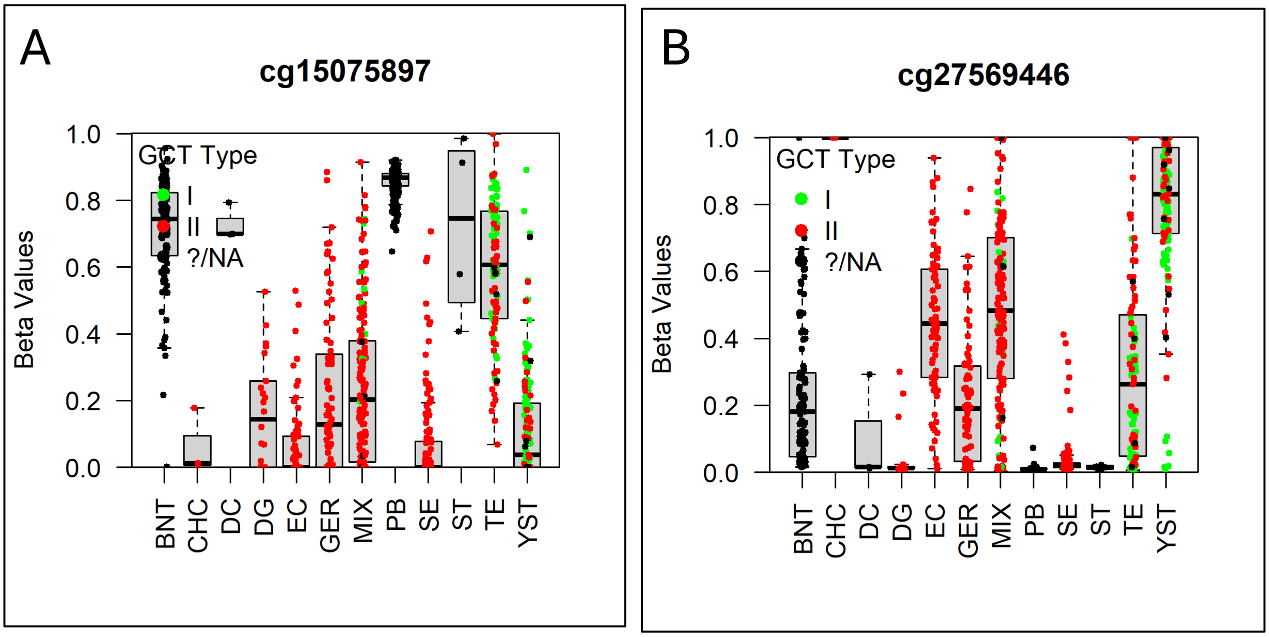


***Figure S14: methylation in two specific probes associated with DPPA3 (A) and RASSF1A (B).*** *Beta-values are plotted for individual samples within the meta-analysis cohort by sample group. Colors indicate GCT type if relevant. cg15075897 is associated with the DPPA3 gene and cg27569446 with RASSF1A. Abbreviations, BNT: benign neighboring testis, CHC: choriocarcinoma, DC: dermoid cyst, DG: dysgerminoma, EC: embryonal carcinoma, GER: germinoma, MIX: GCT with mixed histology, PB: peripheral blood, SE: seminoma, ST: spermatocytic seminoma, TE: teratoma, YST: yolk sac tumor.*

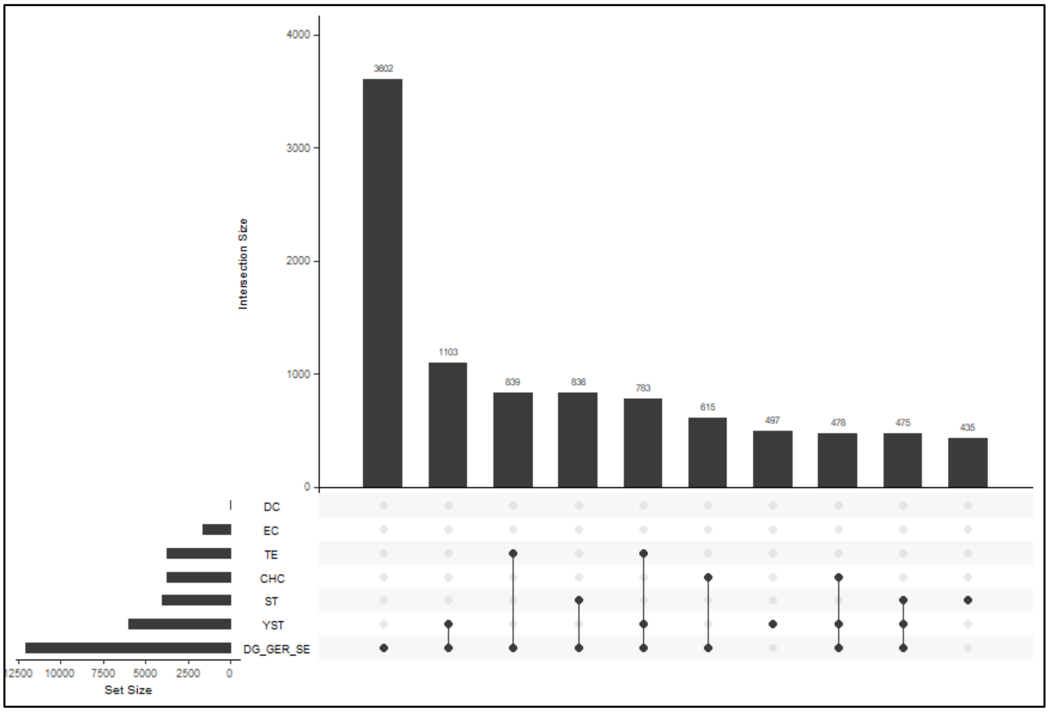


***Figure S15: collectively shared genes derived from differentially methylated regions among GCT histologies.*** *Per sample group, DMR-derived genes were collected (Table S7-13) and overlapping genes were identified using Upset plotting. Intersection Size=total gene number in a single group, or being shared among multiple groups (indicated by bullet points below that column). Set Size=total number of genes per group regardless of presence in other groups. Abbreviations: CHC: choriocarcinoma, DC: dermoid cyst, DG: dysgerminoma, DMR: differentially methylated region, EC: embryonal carcinoma, GER: germinoma, SE: seminoma, ST: spermatocytic tumor, TE: teratoma, YST: yolk sac tumor.*

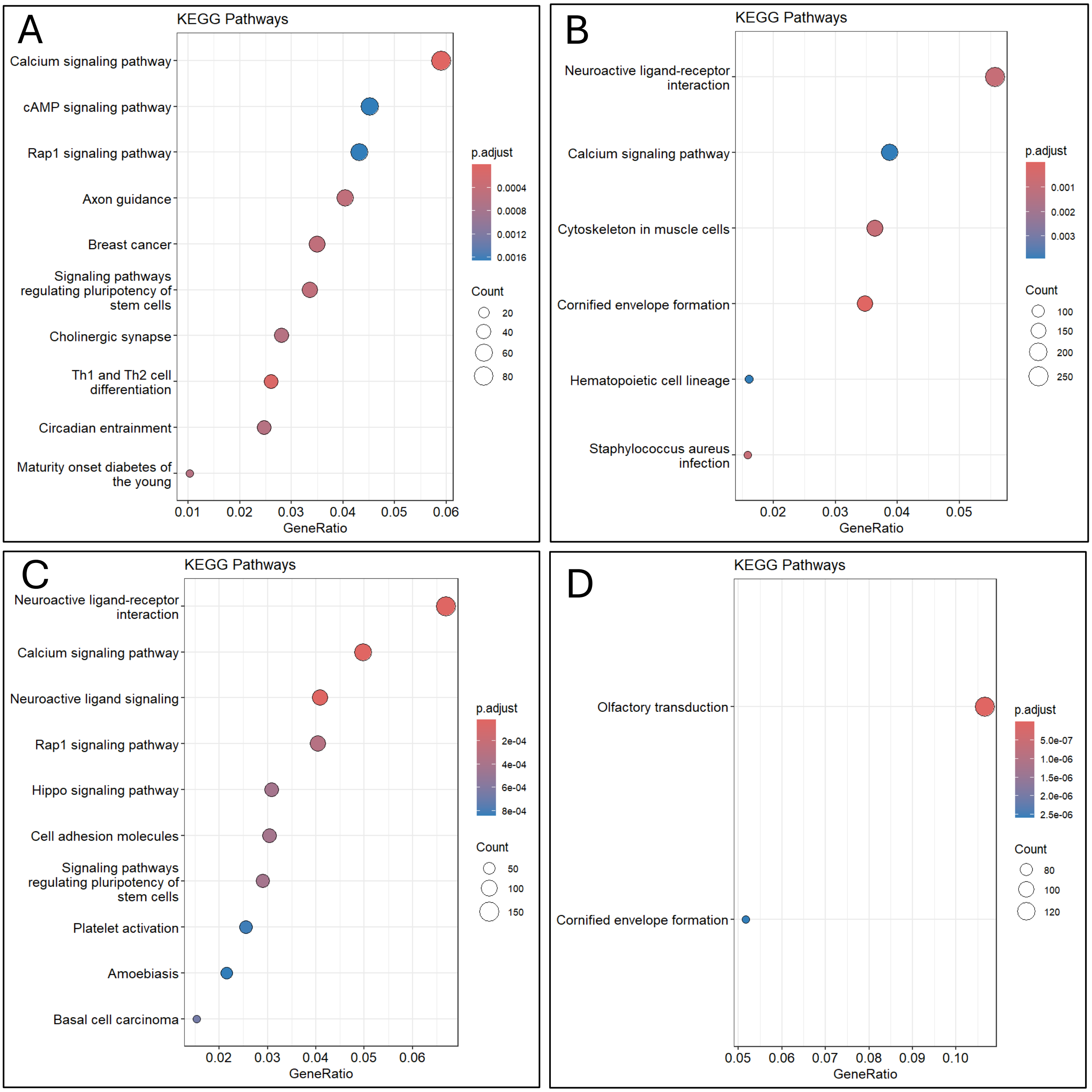
A
KEGG Pathways
Calcium signaling pathway
cAMP signaling pathway
Rap1 signaling pathway
Axon guidance
Breast cancer
Signaling pathways regulating pluripotency of stem cells
Cholinergic synapse
Th1 and Th2 cell differentiation
Circadian entrainment
Maturity onset diabetes of the young
0.01 0.02 0.03 0.04 0.05 0.06
GeneRatio
p.adjust
0.0004
0.0008
0.0012
0.0016
Count
20
40
60
80
B
KEGG Pathways
Neuroactive ligand-receptor interaction
Calcium signaling pathway
Cytoskeleton in muscle cells
Cornified envelope formation
Hematopoietic cell lineage
Staphylococcus aureus infection
0.02 0.03 0.04 0.05
GeneRatio
p.adjust
0.001
0.002
0.003
Count
100
150
200
250
C
KEGG Pathways
Neuroactive ligand-receptor interaction
Calcium signaling pathway
Neuroactive ligand signaling
Rap1 signaling pathway
Hippo signaling pathway
Cell adhesion molecules
Signaling pathways regulating pluripotency of stem cells
Platelet activation
Amoebiasis
Basal cell carcinoma
0.02 0.03 0.04 0.05 0.06
GeneRatio
p.adjust
2e-04
4e-04
6e-04
8e-04
Count
50
100
150
D
KEGG Pathways
Olfactory transduction
Cornified envelope formation
0.05 0.06 0.07 0.08 0.09 0.10
GeneRatio
p.adjust
5.0e-07
1.0e-06
1.5e-06
2.0e-06
2.5e-06
Count
80
100
120

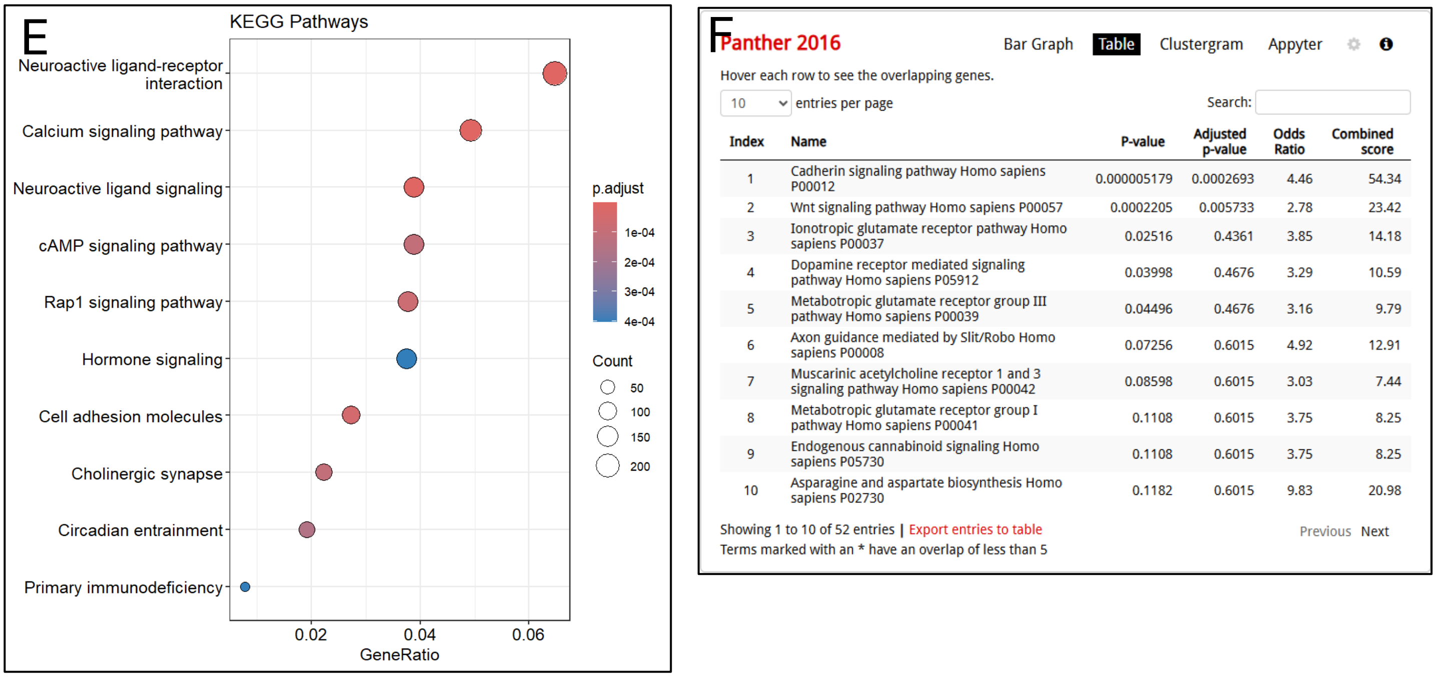


| Index | Name | P-value | Adjusted p-value | Odds Ratio | Combined score |
|---|---|---|---|---|---|
| 1 | Cadherin signaling pathway Homo sapiens P00012 | 0.000005179 | 0.0002693 | 4.46 | 54.34 |
| 2 | Wnt signaling pathway Homo sapiens P00057 | 0.0002205 | 0.005733 | 2.78 | 23.42 |
| 3 | Ionotropic glutamate receptor pathway Homo sapiens P00037 | 0.02516 | 0.4361 | 3.85 | 14.18 |
| 4 | Dopamine receptor mediated signaling pathway Homo sapiens P05912 | 0.03998 | 0.4676 | 3.29 | 10.59 |
| 5 | Metabotropic glutamate receptor group III pathway Homo sapiens P00039 | 0.04496 | 0.4676 | 3.16 | 9.79 |
| 6 | Axon guidance mediated by Slit/Robo Homo sapiens P00008 | 0.07256 | 0.6015 | 4.92 | 12.91 |
| 7 | Muscarinic acetylcholine receptor 1 and 3 signaling pathway Homo sapiens P00042 | 0.08598 | 0.6015 | 3.03 | 7.44 |
| 8 | Metabotropic glutamate receptor group I pathway Homo sapiens P00041 | 0.1108 | 0.6015 | 3.75 | 8.25 |
| 9 | Endogenous cannabinoid signaling Homo sapiens P05730 | 0.1108 | 0.6015 | 3.75 | 8.25 |
| 10 | Asparagine and aspartate biosynthesis Homo sapiens P02730 | 0.1182 | 0.6015 | 9.83 | 20.98 |

Showing 1 to 10 of 52 entries | Export entries to table

Terms marked with an * have an overlap of less than 5

Previous Next

***Figure S16: sample group-specific enriched pathways.*** *Based on DMR-involved genes (Figure S15), significantly enriched KEGG pathways specific to individual sample groups: CHC (A), collective gGCTs (DG, GER, DG) (B), and YST (C) are indicated. Also, pathways for collective gGCTs (D) and collective ngGCTs (CHC, EC, TE, YST) (E) resulting from mutually exclusive genes in both groups are indicated.Per pathway, the number of involved genes, adjusted p-value, and gene ratio are shown. Enriched Panther 2016 pathways are shown (F) for mutually exclusive genes to the YST group. Abbreviations: DMR: differentially methylated region, CHC: choriocarcinoma, DC: dermoid cyst, DG: dysgerminoma, GER: germinoma, SE: seminoma, ST: spermatocytic tumor, TE: teratoma, YST: yolk sac tumor.*

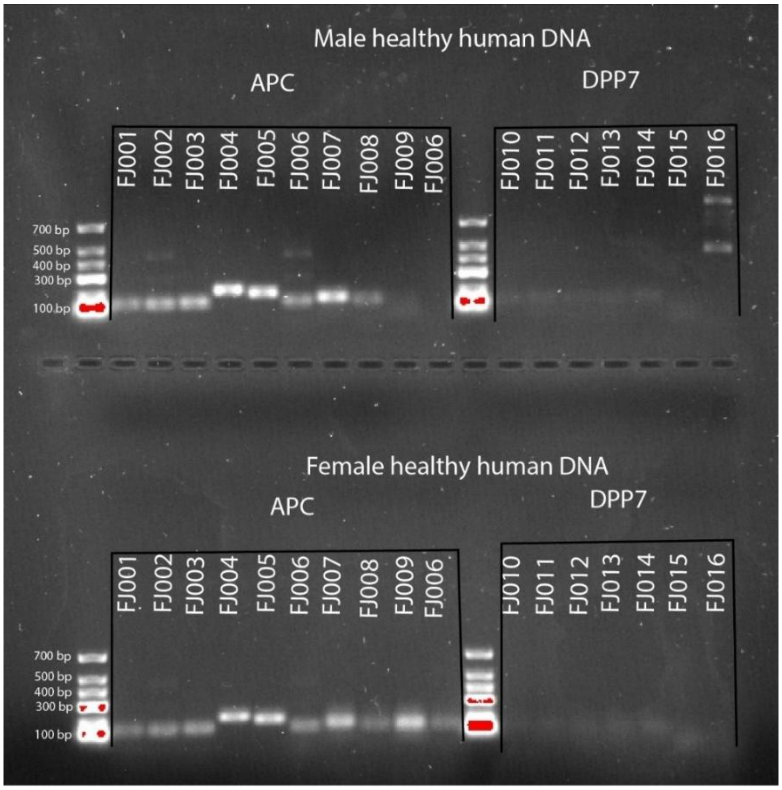


***Figure S17: gel electrophoresis following endpoint PCR for testing different primer combinations covering APC and DPP7 amplicons.*** *16 different primer pairs were tested for APC and DPP7 gene amplification based on healthy male (top) and female (bottom) DNA. Bands indicate target amplification. Amplicons had a size of 65-178 bp.*

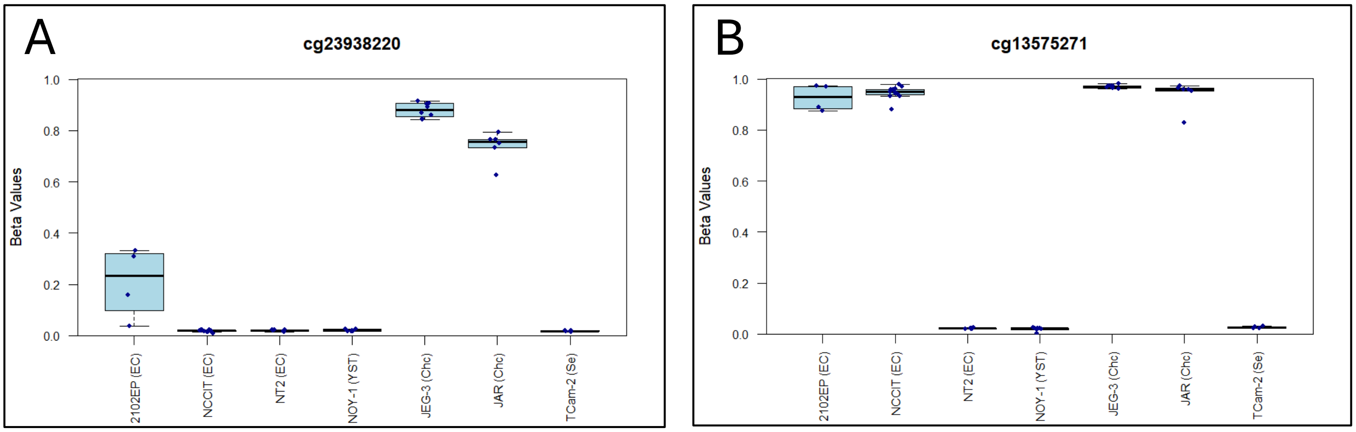


***Figure S18: methylation in two specific probes associated with APC (A) and DPP7 (B) for GCT CLs.*** *Beta-values are plotted for individual samples within the meta-analysis cohort by sample group. cg23938220 is associated with the APC gene and cg13575271 with DPP7. Abbreviations, CHC: choriocarcinoma, EC: embryonal carcinoma, SE: seminoma, YST: yolk sac tumor.*

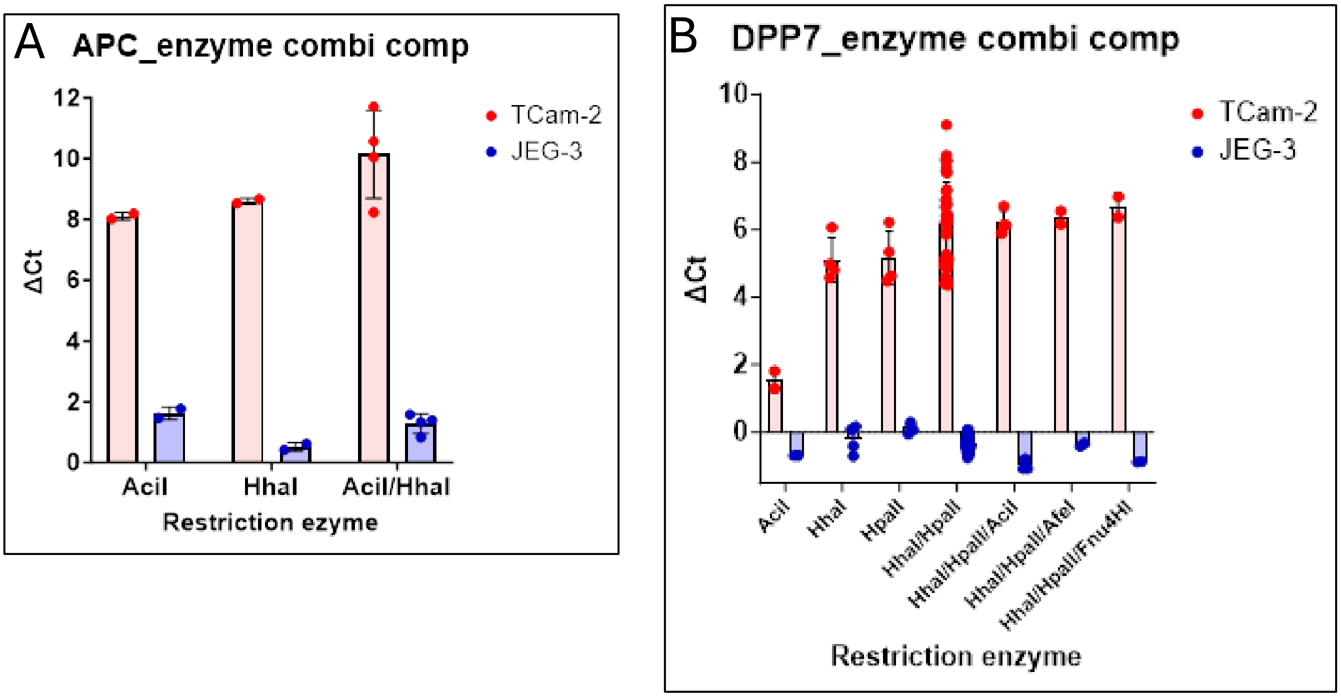


***Figure S19: MSRE digestion efficacy comparison.*** *The effect of MSRE content (x-axis) on digestion efficacy (y-axis) for APC (A) and DPP7 (B) DNA from TCam-2 and JEG-3 GCT cell lines is shown. Error bars indicate standard deviation. Total enzyme input per DNA input varied from 0.02 – 0.49 U/ng.*

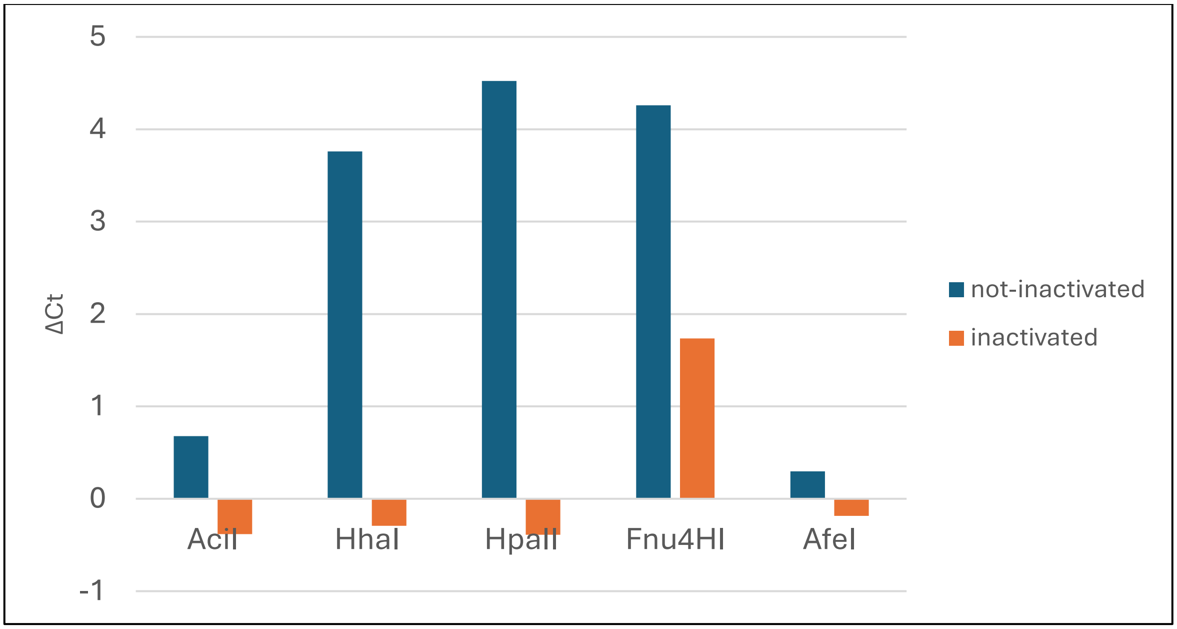


***Figure S20: MSRE digestion efficacy +/- heat inactivation.*** *The effect of MSRE content (x-axis) on digestion efficacy (y-axis) for DPP7 DNA from TCam-2 is shown. For conditions with heat inactivation, enzymes were first incubated at 95°C for 10 minutes. Toal enzyme input was 1 unit with 20 ng DNA (0.05 U/ng). Error bars indicate standard deviation. N=1 per condition.*

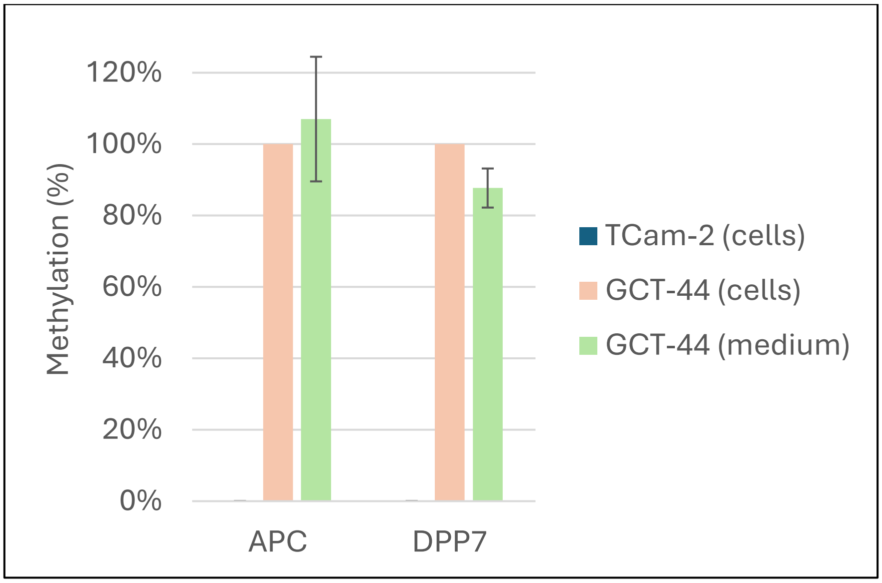


***Figure S21: APC and DPP7 methylation can be measured in conditioned medium of GCT-44 cells.*** *APC and DPP7 methylation was measured using optimized assays within TCam-2 and GCT-44 GCT cell lines, as well as conditioned medium of GCT-44 cells. Results were normalized for methylation % in GCT-44 cells which was set at 100%. N=1 for TCam-2 and GCT-44 cells. N= 2 for GCT-44 (medium) where error bars indicate absolute variation.*

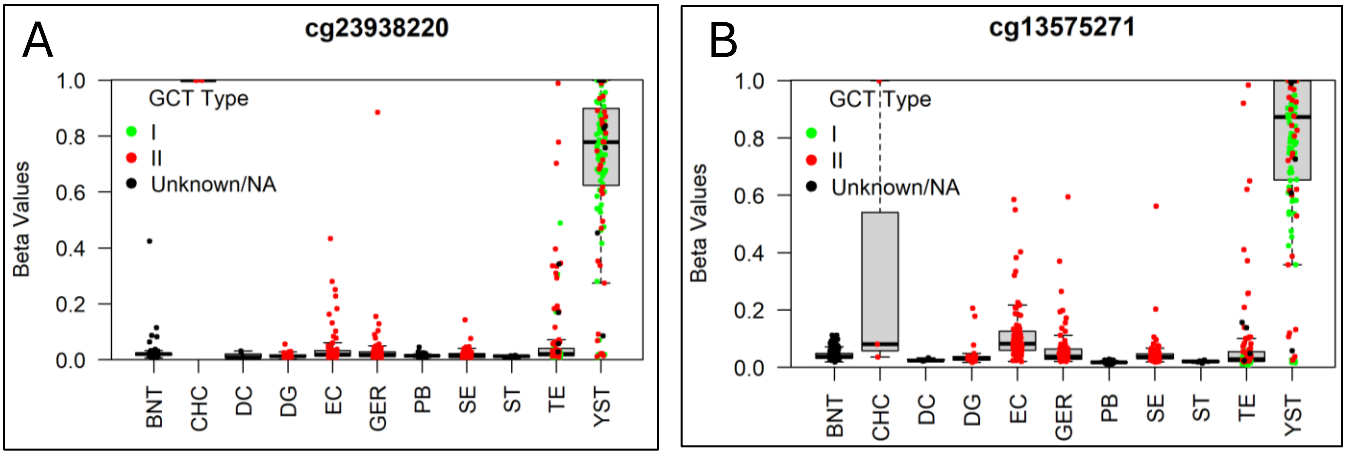


***Figure S22: methylation in two specific probes associated with APC (A) and DPP7 (B).*** *Beta-values are plotted for individual samples within the meta-analysis cohort by sample group. Colors indicate GCT type if relevant. cg23938220 is associated with the APC gene and cg13575271 with DPP7. Abbreviations, BNT: benign neighboring testis, CHC: choriocarcinoma, DC: dermoid cyst, DG: dysgerminoma, EC: embryonal carcinoma, GER: germinoma, PB: peripheral blood, SE: seminoma, ST: spermatocytic seminoma, TE: teratoma, YST: yolk sac tumor.*

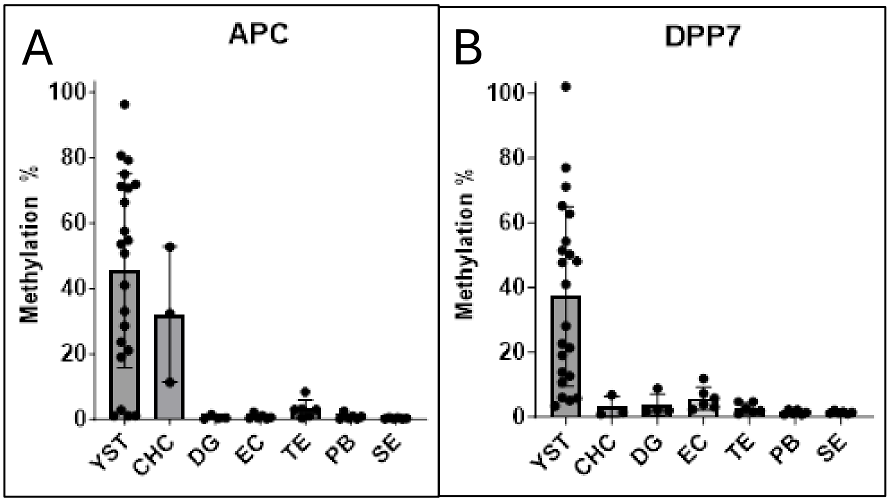


***Figure S23: APC (A) and DPP7 (B) methylation in tumor DNA.*** *Methylation % is plotted for individual samples, per sample group, within the tumor DNA cohort. Error bars indicate standard deviation. Abbreviations, YST: yolk sac tumor, CHC: choriocarcinoma, DG: dysgerminoma, EC: embryonal carcinoma, TE: teratoma, PB: peripheral blood, SE: seminoma.*

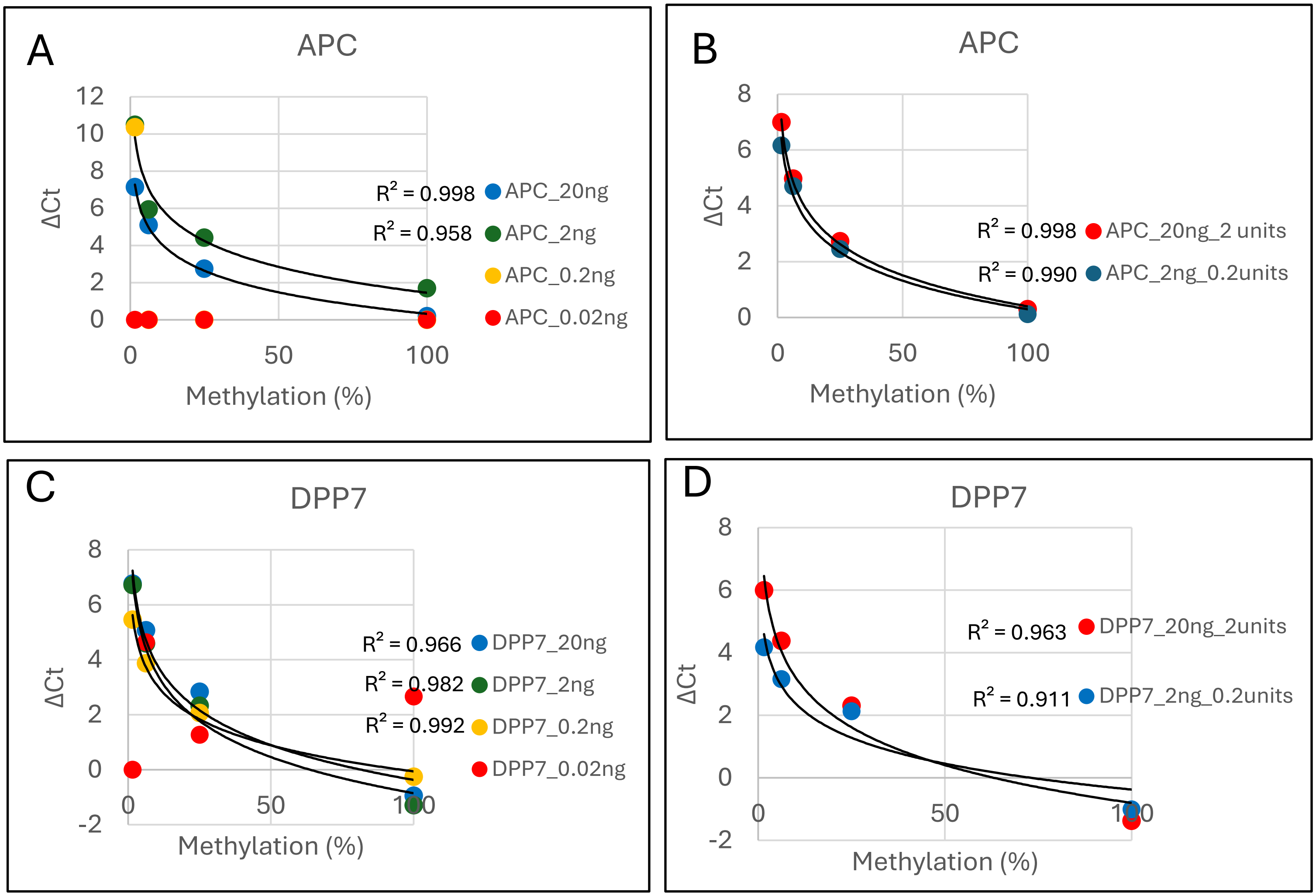


***Figure S24: LOQ experiment for APC and DPP7 methylation assays.*** *For APC (A and B) and DPP7 (C and D), two LOQ experiments were performed. The correlation between ΔCt and methylation % is shown expressed as standard curves, for different DNA and enzyme input conditions. Each sample within a standard curve represents a mix of GCT-44 and NOY-1 (for APC) and JAR and NOY-1 (for DPP7) where GCT-44 and JAR are considered to be 100% methylated and NOY-1 is considered to be 0% methylated. Total enzyme unit input for A and C was 2U. Logarithmic trendlines are fitted (when applicable) through standard curves with correlation coefficients (R2) being indicated. Abbreviations: LOQ: limit of quantification.*

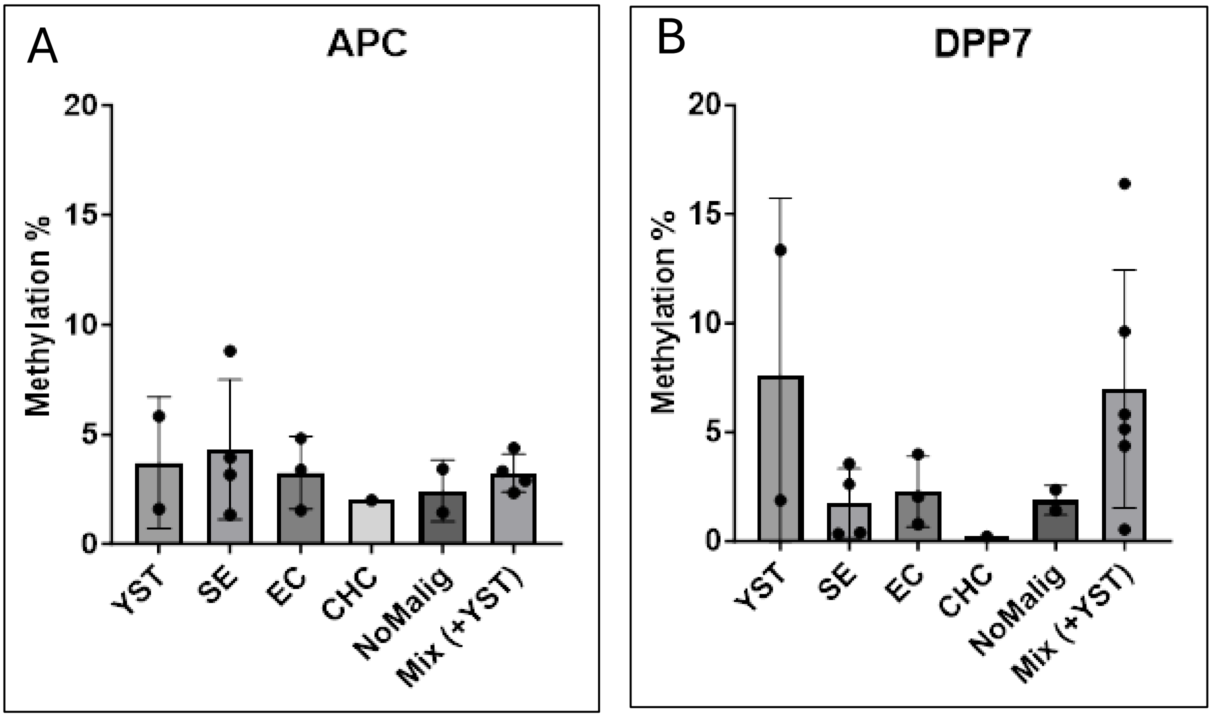


***Figure S25: APC (A) and DPP7 (B) methylation in cfDNA.*** *Methylation % is plotted for individual samples, per sample group, within the liquid biopsy DNA cohort. Error bars indicate standard deviation. Abbreviations, YST: yolk sac tumor, CHC: choriocarcinoma, EC: embryonal carcinoma, SE: seminoma, NoMalig: no malignancy, Mix (+YST): subject has a GCT with multiple histologies including a YST.*

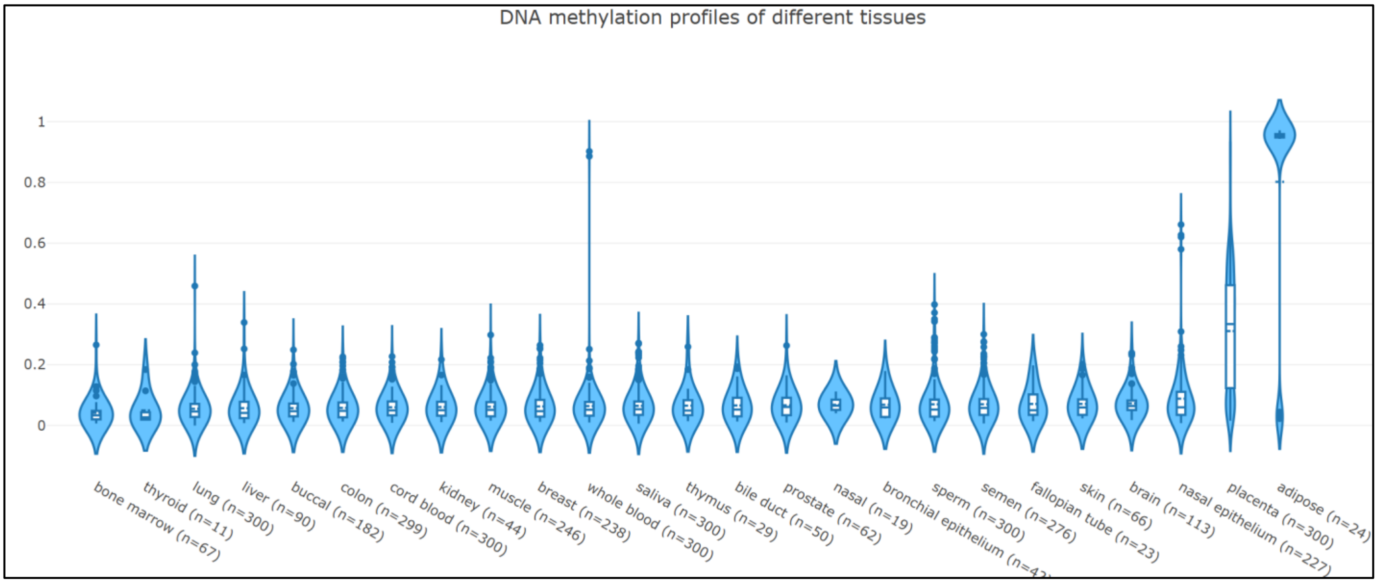


***Figure S26: DNA methylation in different healthy tissues for probe cg13575271.*** *Probe cg13575271 is annotated to gene DPP7. This data) was generated using the EWAS open access platform.* [***https://ngdc.cncb.ac.cn/ewas/datahub***](https://ngdc.cncb.ac.cn/ewas/datahub)

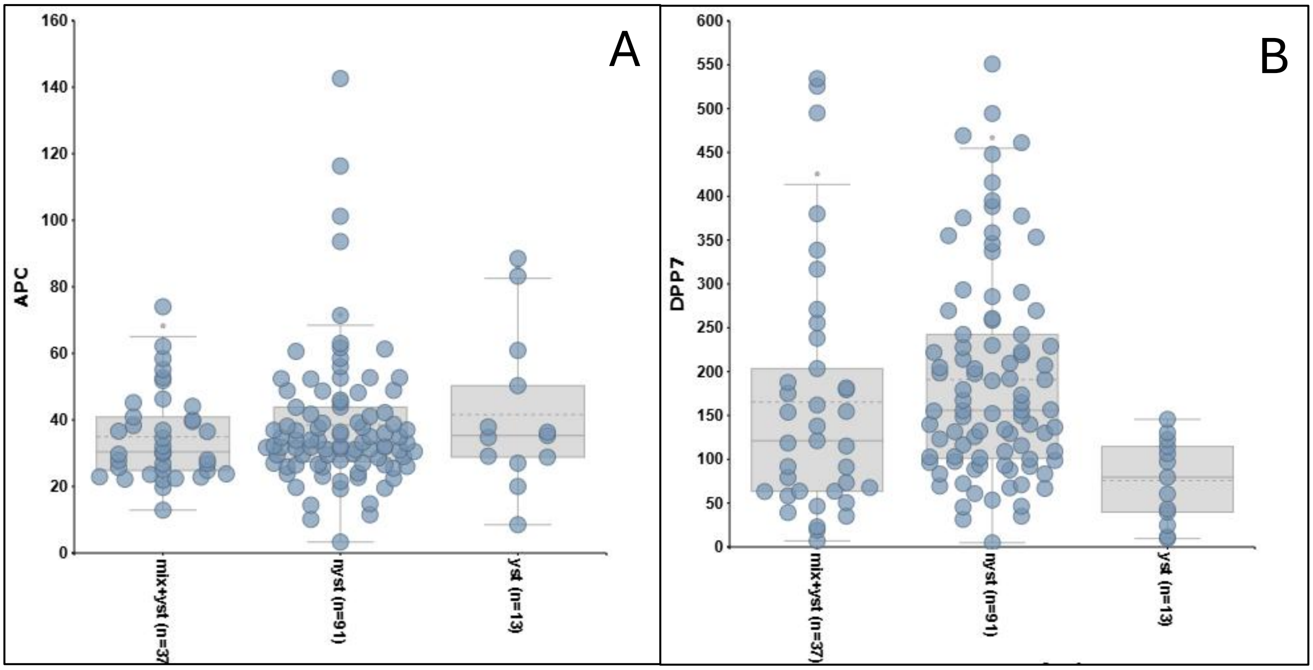


***Figure S27: APC (A) and DPP7 (B) expression in different GCT groups.*** *The nYST group consisted of 4 CHC, 18 EC, 29 MIX-YST, 12 SE, 19 TE, 6 healthy testis, and 3 non-determined samples. The y-axis (gene expression) is linear. Abbreviations: CHC: choriocarcinoma, EC: embryonal carcinoma, YST: yolk sac tumor, SE: seminoma, TE: teratoma, mix+/-YST: mixed GCT with/without a YST component, nYST: non-yolk sac tumor samples. Data is derived from R2 (reference: ps_avgpres_gse3218gse10783ageo141_u133a and ps_avgpres_gse3218gse10783geo141_u133b for APC and DPP7, respectively.*

### Supplementary table legends:

**Table S1: metadata for full meta-analysis cohort.** 1092 samples are included consisting of tissue, cell lines, and healthy peripheral blood. Sample and subject (if relevant)-specific metadata is provided.

**Table S2: metadata for GCT DNA cohort for MSRE-qPCR validation.** 53 samples are included consisting of tissue and healthy peripheral blood. Sample and subject (if relevant)-specific metadata is provided. Abbreviations: GCT: germ cell tumor. MSRE-qPCR: methylation sensitive restriction enzyme-quantitative PCR.

**Table S3: metadata for cfDNA (liquid biopsy) cohort.** 18 samples are included consisting of cfDNA from subjects with a GCT or without malignancy. Sample and subject-specific metadata is provided. Abbreviations: cfDNA: cell-free DNA.

**Table S4: *APC* and *DPP7* primer sequences.** Forward and reverse primer sequences are provided.

**Table S5: unsupervised hierarchical clustering of pure GCT samples.** For each histology, the number of samples is listed, per cluster. Also, the relative sample clustering based on histology and cluster group are shown. Abbreviations: GCT: germ cell tumor.

**Table S6: unsupervised hierarchical clustering of pure + MIX GCT samples.** For each histology, the number of samples is listed, per cluster. Also, the relative sample clustering based on histology and cluster group are shown. Abbreviations: GCT: germ cell tumors, MIX: GCT with multiple histologies.

**Table S7: unsupervised hierarchical clustering of pure GCT + BNT samples.** For each histology, the number of samples is listed, per cluster. Also, the relative sample clustering based on histology and cluster group are shown. Abbreviations: GCT: germ cell tumor, BNT: benign neighbouring testis.

**Table S8: overview of hyper and hypomethylated DMRs per GCT histology.** The mean CpGs number per DMR, mean delta beta-value per DMR, the maximum delta beta-value of a DMR, the mean log10(hmfdr) per DMR, and the maximum log10(hmfdr) per DMR are indicated when comparing a single group versus all other listed groups (including BNT and PB). Abbreviations: GCT: germ cell tumor, DMR: Differentially methylated region, Hmfdr: harmonic mean of the FDR-corrected individual CpG p-values that make up a DMR, BNT: Benign neighbouring testis, PB: Peripheral blood.

**Table S9: DMRs for CHC vs BNT, DC, DG, EC, GER, PB, SE, ST, TE, YST.** DMRs specific for CHC were investigated using the DMRcate algorithm. Each row shows a unique DMR with annotations for associated genetic coordinates, the involved gene and associated annotation/region, multiple statistical metrics, and beta-value differences. The blue color bar length indicates a quantitative scoring between histologies for easy visual comparison per metric. Abbreviations: DMR: differentially methylated region, CHC: choriocarcinoma, BNT: benign neighbouring testis, DC: dermoid cyst, DG: dysgerminoma, EC: embryonal carcinoma, GER: germinoma, PB: peripheral blood, SE: seminoma, ST: spermatocytic tumor, TE: teratoma, YST: yolk sac tumor.

**Table S10: DMRs for DC vs BNT, CHC, DG, EC, GER, PB, SE, ST, TE, YST.** DMRs specific for DC were investigated using the DMRcate algorithm. Each row shows a unique DMR with annotations for associated genetic coordinates, the involved gene and associated annotation/region, multiple statistical metrics, and beta-value differences. See Table S9 for abbreviations.

**Table S11: DMRs for EC vs BNT, CHC, DC, DG, GER, PB, SE, ST, TE, YST.** DMRs specific for EC were investigated using the DMRcate algorithm. Each row shows a unique DMR with annotations for associated genetic coordinates, the involved gene and associated annotation/region, multiple statistical metrics, and beta-value differences. See Table S9 for abbreviations.

**Table S12: DMRs for ST vs BNT, CHC, DC, DG, EC, GER, PB, SE, TE, YST**. DMRs specific for ST were investigated using the DMRcate algorithm. Each row shows a unique DMR with annotations for associated genetic coordinates, the involved gene and associated annotation/region, multiple statistical metrics, and beta-value differences. See Table S9 for abbreviations.

**Table S13: DMRs for TE vs BNT, CHC, DC, DG, EC, GER, PB, SE, ST, YST**. DMRs specific for TE were investigated using the DMRcate algorithm. Each row shows a unique DMR with annotations for associated genetic coordinates, the involved gene and associated annotation/region, multiple statistical metrics, and beta-value differences. See Table S9 for abbreviations.

**Table S14: DMRs for YST vs BNT, CHC, DC, DG, EC, GER, PB, SE, ST, TE**. DMRs specific for YST were investigated using the DMRcate algorithm. Each row shows a unique DMR with annotations for associated genetic coordinates, the involved gene and associated annotation/region, multiple statistical metrics, and beta-value differences. See Table S9 for abbreviations.

**Table S15: DMRs for DG, GER, SE vs BNT, CHC, DC, EC, PB, ST, TE, YST**. DMRs specific for gGCTs were investigated using the DMRcate algorithm. Each row shows a unique DMR with annotations for associated genetic coordinates, the involved gene and associated annotation/region, multiple statistical metrics, and beta-value differences. See Table S9 for abbreviations.

**Table S16: DMRs for CHC, DC, EC, ST, TE, YST vs BNT, DG, GER, PB, SE**. DMRs specific for ngGCTs (+DC and ST) were investigated using the DMRcate algorithm. Each row shows a unique DMR with annotations for associated genetic coordinates, the involved gene and associated annotation/region, multiple statistical metrics, and beta-value differences. See Table S9 for abbreviations.

**Table S17: DMRs for CHC, DC, DG, EC, GER, MIX, SE, ST, TE, YST vs BNT, PB**. DMRs specific for all GCTs were investigated using the DMRcate algorithm. Each row shows a unique DMR with annotations for associated genetic coordinates, the involved gene and associated genomic feature annotation, multiple statistical metrics, and beta-value differences. See Table S9 for abbreviations.

**Table S18: Genes within DMRs per subgroup and their genomic feature annotations**. Genes involved in different DMR-analyses (Table S9-17) are listed, including the gene proportion unique to that subgroup, as well as corresponding genomic feature annotations.

**Table S19: Full list of DMR-derived genes exclusive per subgroup**. Genes involved in different DMR-analyses (Table S9-17), exclusive to each subgroup, are listed. The total number of genes per subgroup is the same as indicated in Table S18 for the “number of genes unique in that group” row.
